# Chromosomes function as a barrier to mitotic spindle bipolarity in polyploid cells

**DOI:** 10.1101/572099

**Authors:** Alix Goupil, Maddalena Nano, Gaëlle Letort, Delphine Gogendeau, Carole Pennetier, Renata Basto

**Author notes:** These authors contribute equally to this work (Alix Goupil, Maddalena Nano).

## Abstract

Whole genome duplications (WGDs) are found in a variety of tumors and are associated with chromosomal instability (CIN) and poor prognosis [1,2]. When induced experimentally, through cytokinesis failure, polyploid cells generate tumors [3]. Cytokinesis failure results in the accumulation of double DNA content, but also of cytoplasmic organelles, such as centrosomes, which are the major microtubule (MT) organizing centers of animal cells. Importantly, even if there is a correlation between polyploidy and CIN [4], the underlying mechanisms generating error-prone mitosis in cells with extra DNA and extra centrosomes are not known. When considering polyploid mitosis, it is essential to take into account the increase in MT nucleation due to the presence of extra centrosomes and extra DNA. The presence of supernumerary centrosomes in a cell, centrosome amplification [5], is associated with mitotic spindle multipolarity and CIN [6–9]. Importantly, additional MTs can be nucleated from the chromatin (chromatin mediated pathway-CMP) or from pre-existing MTs-through the Augmin pathway. We hypothesized that the increase in DNA and centrosome content in a cell could lead to an increased MT mass, which might account for abnormal mitosis described in polyploid cells [4, 10, 11, 12]. Using genetics, live imaging and modeling approaches, we investigated the mechanisms establishing multipolarity *in vivo* in polyploid cells. We found that MT nucleation from the centrosomes is the major contributor to multipolarity, while other pathways seem to play minor roles. Unexpectedly, we found that even if Ncd/HSET, plays an essential role in promoting centrosome clustering in early mitosis, the increase in chromosome mass associated with cytokinesis failure functions as a barrier to centrosome clustering into two main poles. Our work provides a mechanistic link between polyploidy and the generation of CIN.

## Results and Discussion

### Polyploid mitosis are multipolar

In order to characterize cell division in polyploid neural stem cells (NSCs) of *Drosophila* larval brain, also called neuroblasts (NBs) we used genetic conditions that induce cytokinesis failure. As described in [13], the hypomorphic mutation in the spaghetti squash (*sqh*) gene, which encodes the regulatory light chain of myosin II [14] results in the generation of polyploid NBs that divide in a multipolar way, in contrast to diploid control (Ctrl) NBs (Figure 1A-B). Moreover, to precisely control the generation of polyploidy in a different loss-of-function context, we used the knock down (KD) of Pavarotti (Pav), the human homologue of MKLP1 [15], through inducible RNA interference (RNAi) (see methods). As we observed in *sqh*^*mut*^ NBs [13], analysis of Pav^KD^ third instar brains revealed the presence of large NBs containing large amounts of DNA, when compared to diploid controls (Ctrl) (Supplementary Figure 1A). Interestingly, Pav^KD^ NBs displayed different sizes and contained increased centrosome numbers (Supplementary Figure 1B), revealing different degrees of polyploidy.

**Figure 1:**
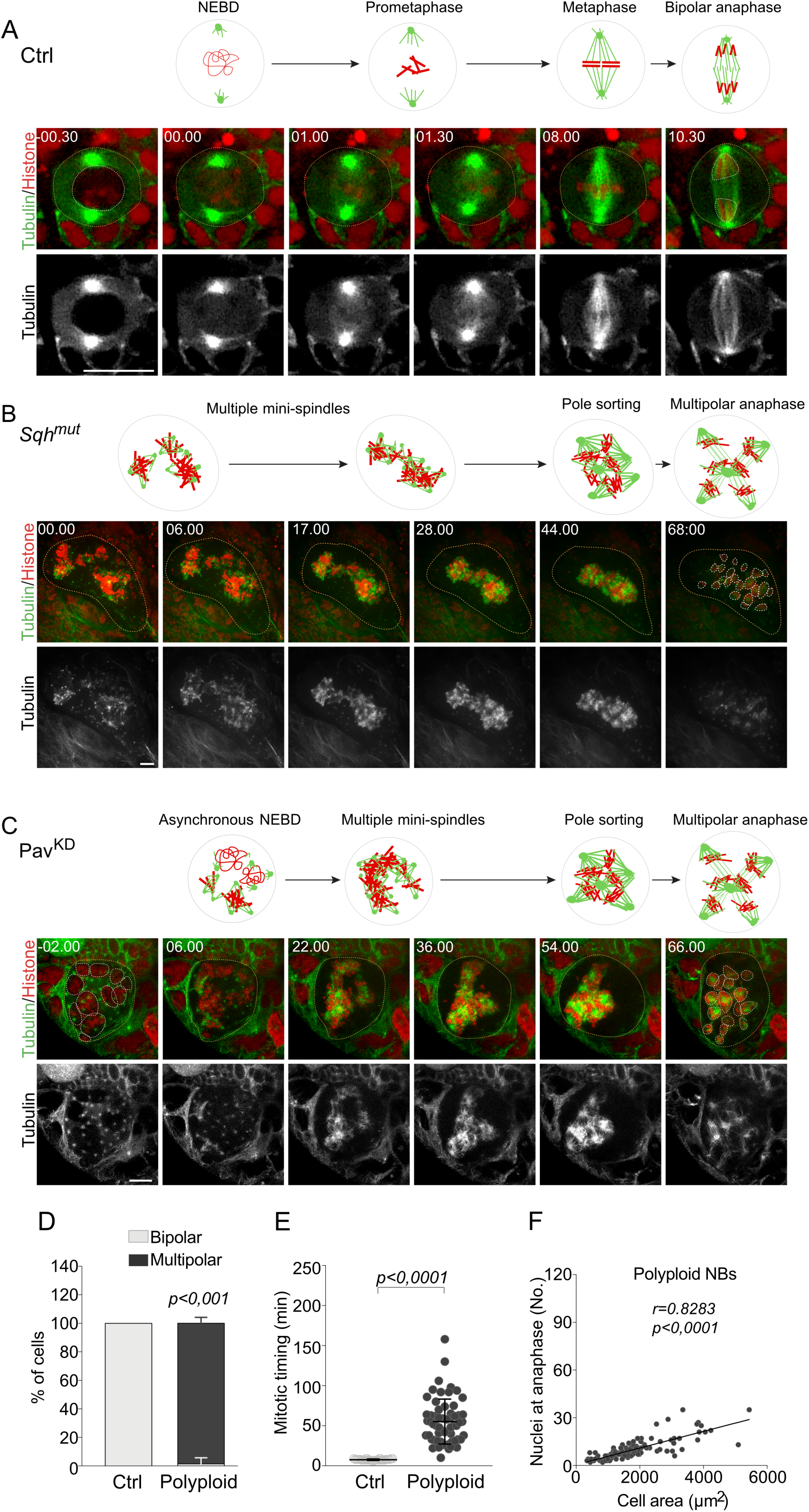
Polyploid cells undergo multipolar mitosis. (A-C) Stills of time-lapse movies of mitotic neuroblasts (NBs) expressing Tubulin-GFP (in green and grey in the bottom insets) and Histone 2B-RFP (in red). Orange and white dotted circles surround cells and nuclei, respectively. Time of mitosis is indicated in minutes(min).seconds(sec). Time 00.00 corresponds to nuclear envelope breakdown (NEBD) for (A and B) and to the beginning of time-lapse acquisition for (C). Schematic representations of mitosis are shown above the stills. (A) Diploid control NB with two centrosomes assembles a bipolar spindle and divides asymmetrically to give rise to two daughter cells. (B and C) Polyploid *sqh*^*mut*^ (B) and Pav^KD^ (C) NBs present several nuclei that condense and assemble in a complex structure centered in the cells. An initial clustering decreases the number of MT-nucleating sites, shown as green dots, allowing a switch from multiple mini-spindles to a unique highly multipolar spindle. At anaphase onset, a high number of nuclei are generated. (D) Graph bars showing the percentage of cells presenting bipolar and multipolar mitosis in Ctrl (n=34 cells from 2 brains) and polyploid (Pav^KD^) NBs (n=107 cells from 37 brains). Statistical significance was determined using a Multiple t-test. (E) Dot plot showing the time spent in mitosis for Ctrl (n=34 cells from 2 brains) and polyploid NBs (n=60 cells from 31 brains). Statistical significance was determined using a Mann-Whitney test. (F) XY plot and corresponding linear regression between the number of nuclei generated at anaphase and polyploid cell area (n=107 cells from 37 brains). Statistical significance of the correlation was determined by a Spearman r test. R corresponding to the correlation coefficient. P corresponds to the p-value and lines represent the mean ± SD. Scale bar = 10µm.

To follow mitosis by time-lapse microscopy, we used fly lines expressing transgenes encoding α-Tubulin tagged with GFP (Tubulin-GFP) and Histone2B tagged with RFP (Histone-RFP) to monitor spindle microtubules (MTs) and chromosomes, respectively. Ctrl diploid NBs divided asymmetrically, as described previously [16, 17]. As cells entered mitosis, centrosomes formed two robust MT asters, assembled a bipolar spindle within just a few minutes and divided in a bipolar manner to generate two daughter cells (Figure 1A, 1D and Video 1). In *sqh*^*mut*^ or Pav^KD^ NBs, several active nucleating centrosomes could be identified, consistent with the fact that cytokinesis failure leads to the accumulation of extra centrosomes (Figure 1B-C, Supplementary Figure 1B and Videos 2 and 3). In both conditions, mitoses were mostly multipolar (Figure 1B-D).

Careful analysis of polyploid NB divisions confirmed our initial observations. As described in *sqh*^*mut*^ NBs [13], Pav^KD^ NB presented several nuclei that asynchronously entered in mitosis. As asynchronous NEBD occurred, extra centrosomes could be seen clustering. During this process, multiple centrosomes gathered in more than two groups, while chromosomes condensed and assembled in a structure frequently centered within the cytoplasm. These multipolar spindles were maintained in a multipolar configuration for a certain period of time before undergoing multipolar anaphase, generating several nuclei (Figure 1C-D and Movie 3). As we decided to use Pav^KD^ to induce polyploidy, we will refer to Pav^KD^ NBs simply as polyploid NBs.

Next, we characterized different parameters of polyploid mitosis. First, we analyzed mitotic duration in polyploid NBs, here defined as the time elapsed between NEBD and anaphase onset. We found that the large majority of polyploid cells took more time to divide than diploid controls (55.12±28.07 min in polyploid vs 7.37±0.53 min in Ctrl, p<0,0001, Figure 1E, Supplementary Table 1). Interestingly, this increased mitotic duration did not necessarily correlate with the degree of polyploidy (as inferred by cell size). Indeed, small polyploid cells could take longer than larger cells to divide (Supplementary Figure 1C-D and Video 4). The increase in mitotic timing is a known feature of cells with extra centrosomes [7, 18], where it reflects a delay in turning off the spindle assembly checkpoint (SAC), which monitors kinetochore-MT attachments [19]. These observations suggest that in polyploid cells achieving accurate kinetochore-MT attachment takes longer than in diploid cells, independently of the degree of polyploidy.

Second, we wanted to assess the outcome of mitosis. While canonical cell division normally involves the separation of one nucleus in two, polyploid mitosis frequently began with- and generated-multiple nuclei. To quantitatively assess the outcome of mitosis, we defined a parameter that we called nuclear index (NI), calculated as the ratio between the number of nuclei formed at anaphase (see methods for detailed analysis) over the number of nuclei at mitotic entry (Supplementary Figure 1E). Cells were categorized in three groups: (a) NI inferior or equal to 1; (b) NI between 1 and 2 and (c) NI superior or equal to 2 (Supplementary Figure 1E-F). Ctrl diploid NBs always presented a NI of 2 (2 post-mitotic nuclei/1 pre-mitotic nucleus = 2). It is important to note that a nuclear index of 2 can be obtained in any condition where the number of post-mitotic nuclei is double the number of pre-mitotic nuclei. The vast majority of polyploid NBs presented a NI higher than 1 (Categories (b) + (c), Supplementary Figure 1F), indicating that at each cycle, the number of nuclei generated at anaphase is higher than the number of nuclei at mitotic entry. Importantly, almost half of the NBs displayed a NI higher or equal to 2 (Cat. (c), Supplementary Figure 1F), which resulted from more than the doubling of the number of nuclei at mitotic exit, albeit not arisen from one nucleus dividing into two. Interestingly, a positive correlation between the number of nuclei at mitotic exit and cell area was noticed (Figure 1F). Very large cells (>6000 μm^2^), which contained more centrosomes and chromosomes than smaller polyploid cells, always generated a higher number of nuclei at mitotic exit when compared to the number of nuclei at mitotic entry (data not shown). These results suggest that the degree of polyploidy impacts the outcome of division, with more and more nuclei generated at each cycle.

Altogether, our results show that polyploidy induced through cytokinesis failure *in vivo*, is associated with an inherent tendency to lose bipolarity. Further, they show that larger cells, which contain more DNA and more centrosomes, generate more nuclei at mitotic exit than smaller cells.

### Investigating the contribution of MT nucleation pathways to multipolar mitosis in polyploid NBs

At least three pathways contribute to MT nucleation during mitosis in animal cells (Figure 2A) [20, 21]. We hypothesized that the multipolar outcome typical of polyploid NBs mitosis could result from an excess of MTs nucleated from extra centrosomes, chromosomes and pre-existing MTs. We tested their relative contribution by decreasing the expression of key players in each pathway individually.

**Figure 2:**
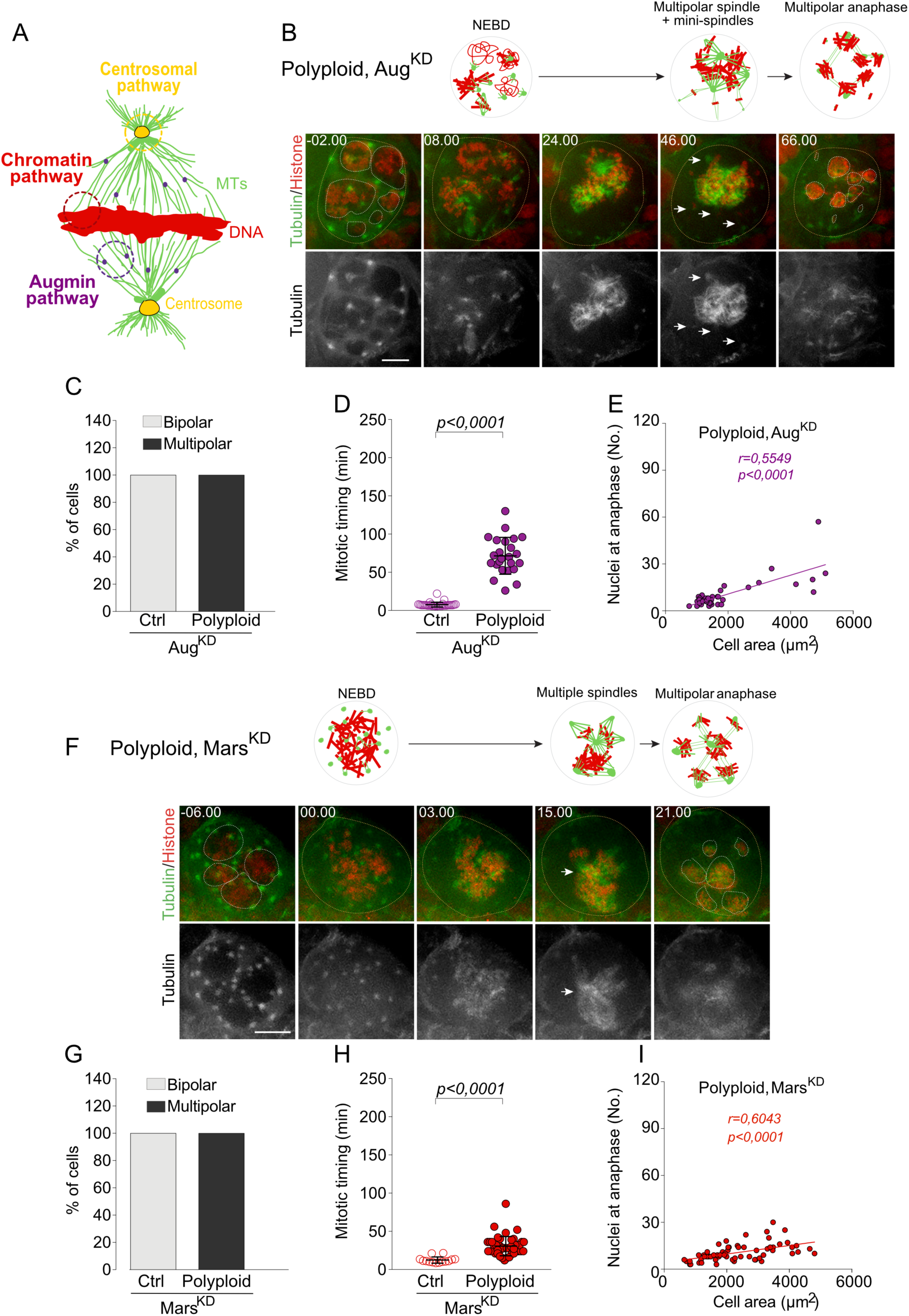
MT-nucleation from the Augmin and chromatin-mediated pathways contribute to the formation of a unique multipolar spindle. (A) Schematic representation of a bipolar spindle. MTs nucleate through three different pathways: the centrosome (in yellow), the chromatin (in red) and from pre-existing MTs, using the Augmin pathway (in purple). DNA and MTs are represented in red and green, respectively. (B) Stills of time-lapse movies of mitotic polyploid, Aug^KD^ NBs expressing Tubulin-GFP (in green and grey in the bottom insets) and Histone 2B-RFP (in red). (C) Graph bars showing the percentage of cells presenting bipolar and multipolar mitosis in Ctrl, Aug^KD^ (n=30 cells from 3 brains) and polyploid, Aug^KD^ NBs (n=45 cells from 15 brains). (D) Dot plot showing the time spent in mitosis for Ctrl, Aug^KD^ (n=30 cells from 3 brains) and polyploid, Aug^KD^ NBs (n=24 cells from 10 brains). (E) XY plot and corresponding linear regression between the number of nuclei generated at anaphase and polyploid, Aug^KD^ cell area (n=45 cells from 15 brains). (F) Stills of time-lapse movies of mitotic polyploid, Mars^KD^ NBs expressing Tubulin-GFP (in green and grey in the bottom insets) and Histone 2B-RFP (in red). (G) Graph bars showing the percentage of cells presenting bipolar and multipolar mitosis in Ctrl, Mars^KD^ (n=13 cells from 1 brain) and polyploid, Mars^KD^ NBs (n=63 cells from 36 brain). (H) Dot plot showing the time spent in mitosis for Ctrl, Mars^KD^ (n=13 cells from 1 brain) and polyploid, Mars^KD^ NBs (n=46 cells from 28 brains). (I) XY plot and corresponding linear regression between the number of nuclei generated at anaphase and polyploid, Mars^KD^ cell area (n=63 cells from 36 brains). (B and F) orange and white dotted circles surround cells and nuclei, respectively. Time of mitosis is represented in min.sec and time 00.00 corresponds to NEBD. Schematic representations of mitosis are shown above the stills. (D and H) Statistical significance was determined using a Mann-Whitney test and lines represent the mean ± SD. (E and I) Statistical significance of the correlation was determined by a Spearman r test. R corresponding to the correlation coefficient. P corresponds to the p-value. Scale bar = 10µm.

First, we interfered with the Augmin pathway, which promotes branched MT nucleation from pre-existing MTs [22, 23]. We used a previously validated RNAi system to knock down (KD) the Dgt2 subunit of the Augmin complex, which was shown to decrease Augmin activity (referred to as Aug^KD^) [20, 23]. Diploid, Aug^KD^ NBs did not display any defect in cell division (Supplementary Figure 2A and Video 5). However, polyploid, Aug^KD^ NBs assembled multipolar spindles and nuclear divisions were always multipolar (Figure 2B-C and Video 6). The initial step of centrosome clustering occurred similarly to what observed in polyploid NBs. However, in certain NBs, we noticed the presence of mini-spindles (Figure 2B, time 46.00min-white arrows and depicted in scheme above). This suggests that, in polyploid cells, Aug contributes to the assembly of a single spindle structure, even if multipolar (Figure 2B). Mitotic duration was also substantially increased in polyploid, Aug^KD^ NBs (71.30±24.08 min in polyploid, Aug^KD^ vs 7.38±3.07 min in diploid, Aug^KD^, p<0,0001), even when compared to polyploid NBs (55.12±28.07 min, p<0,0075) (Figure 2D and Supplementary Table 1). These results show that Aug depletion in polyploid cells delays mitotic progression. The mitotic delay induced by Aug^KD^ is likely due to its role in influencing MT directionality [24], which might impact the overall organization of the spindle and thus, the proper capture of kinetochores and SAC satisfaction.

Next, we interfered with the chromatin mediated pathway (CMP). We depleted Mars, the homologue of hepatoma up-regulated protein (Hurp), a chromatin-associated spindle assembly factor, using tools previously used in *Drosophila* [20, 25]. A reduction in CMP did not reduce multipolarity, as polyploid, Mars^KD^ NBs always divided in a multipolar fashion (Figure 2F-G and Video 7), while diploid, Mars^KD^ NBs always divided in a bipolar manner (Supplementary Figure 2B, Figure 2G and Video 8). As with Aug depletion, polyploid, Mars^KD^ NBs presented defects in spindle formation. Indeed, the majority of polyploid, Mars^KD^ cells, assembled several individual spindles and - in certain occasions - these seemed to share a spindle pole (Figure 2F-white arrow). Interestingly, the decrease in CMP reduced mitotic timing compared to unperturbed polyploid NBs (30.34±12.32 min in polyploid, Mars^KD^ vs 55.12 ± 28.07 min in polyploid NBs, p<0.0001) (Figure 2H and Supplementary Table 1), indicating that the decrease in CMP-dependent MT nucleation favors mitotic progression. In this case, the reduction in MT density generated by CMP most likely favors kinetochore-MT attachment and so anaphase onset.

Interestingly, while the correlation coefficient *r* was lower in polyploid, Aug^KD^ and polyploid, Mars^KD^ NBs, when compared to polyploid NBs, we still observed a positive correlation between the number of nuclei at anaphase and NB area in these genetic combinations (Figure 2E and 2I, p<0,0001 for both). Importantly, Mars depletion, but not Aug, resulted in a lower regression slope (p=0,0002) indicative of a reduction in the number of nuclei generated at anaphase for a given cell size and so polyploid level (Figure 2I and Supplementary Table 1).

These results suggest that even if neither a reduction in the Augmin or CM pathways can rescue the multipolarity typical of polyploid NBs, the reduction in the CMP seems to limit the number of daughter nuclei, suggesting a reduction in the extend of multipolarity. Further, these pathways seem to play important roles in maintaining the extra DNA and extra centrosomes of polyploid NBs in a unique (even if multipolar) structure.

### Centrosomes greatly contribute to multipolarity in polyploid NBs

We then tested the contribution of the centrosomal pathway to multipolar spindle assembly. The centrosome is the major MT organizing center of animal cells [26, 27]. The presence of centrosomes, albeit non-essential for spindle assembly in *Drosophila* NBs, ensures faster MT nucleation and spindle robustness [28, 29]. To reduce centrosome-dependent MT-nucleation, we used a mutation in the *sas-4* gene (CPAP in humans), which encodes an essential centriole duplication gene [28, 30]. Surprisingly, small to medium size polyploid, *sas4*^*mut*^ NBs started by assembling an amorphous MT-structure around the DNA, which evolved into an almost perfect bipolar spindle, albeit larger than in diploid, *sas4*^*mut*^ NBs (Figure 3A, 3C, Supplementary Figure 2C and Videos 9 and 10). The ability of centrosome loss to sustain bipolar spindle assembly was striking, but limited by cell size, as large polyploid, *sas4*^*mut*^ NBs divided in a multipolar manner even in the absence of centrosomes or when centrosome number was highly reduced (Figure 3B-C and Video 11). Mitotic timing was decreased in polyploid, *sas4*^*mut*^ NBs when compared to polyploid NBs (38.19±14.41 min in polyploid*, sas4*^*mut*^ vs 55.12 ± 28.07 min in polyploid NBs, p=0,0073) (Figure 3D and Supplementary Table 1), supporting the view that increased MT nucleation found in polyploid NBs delays inactivation of the SAC. Consistent with an increase in bipolarity, the NI was below 2 in the large majority of polyploid, *sas4*^*mut*^ NBs and significantly decreased when compared to polyploid NBs (1.24 ± 0.42 vs 1.86±0.64, p<0.0001) (Supplementary Figure 1F and Supplementary Table 1). Indeed, this was the only condition where the NI was significantly reduced. It is important to mention that the number of nuclei at anaphase still correlated with cell area, though the regression slope was decreased when compared to polyploid cells (Figure 3E and Supplementary Table 1, p<0,0001), since more than two nuclei were only generated in medium-large size polyploid, *sas4*^*mut*^ NBs.

**Figure 3:**
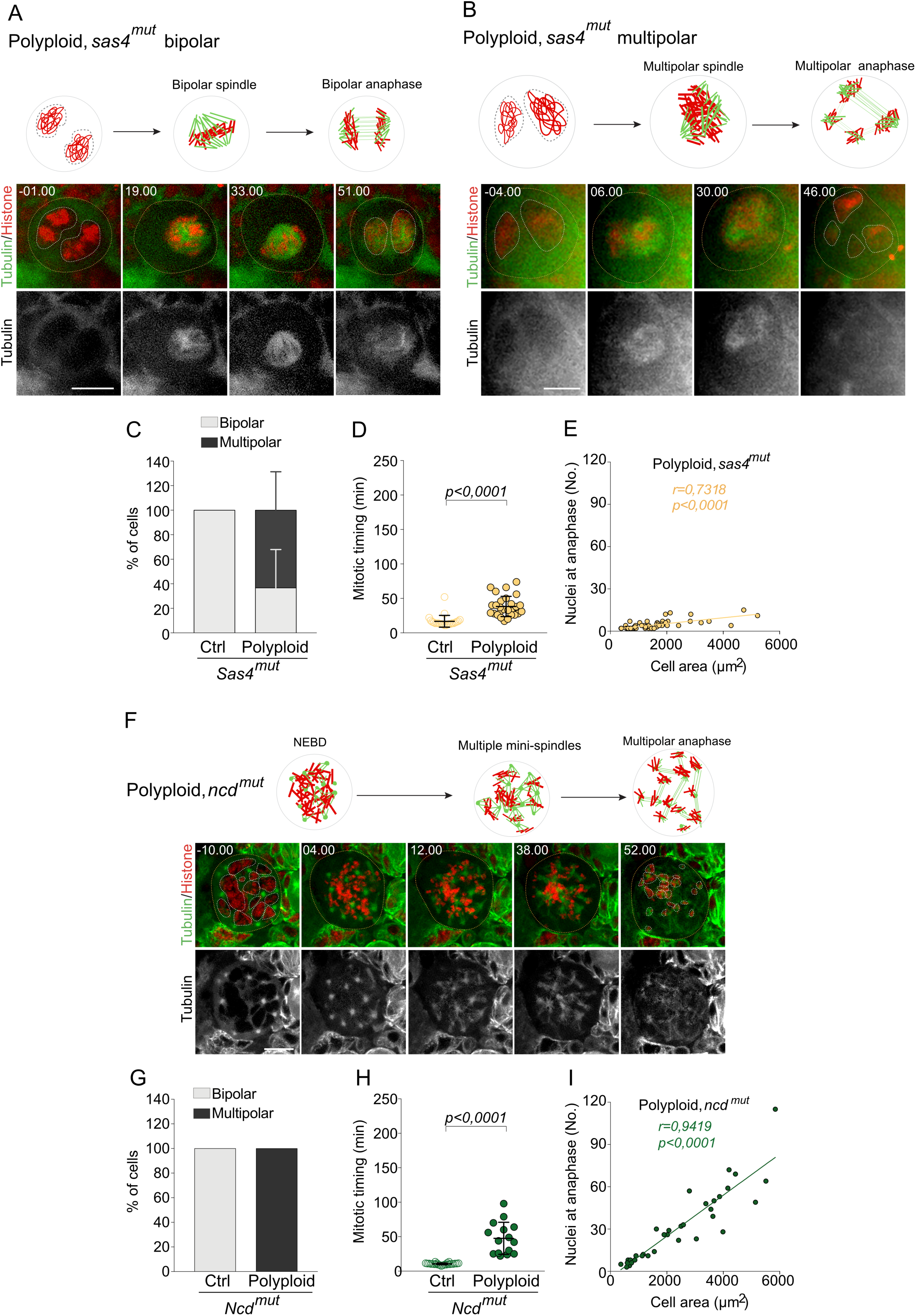
Centrosomes are major contributors to multipolarity in polyploid cells, but they initially cluster in a Ncd-dependent fashion. (A-B) Stills of time-lapse movies of polyploid, *sas4*^*mut*^ NBs expressing Tubulin-GFP (in green and grey in the bottom insets) and Histone 2B-RFP (in red). (C) Graph bars showing the percentage of cells presenting bipolar and multipolar mitosis in Ctrl, *sas4*^*mut*^ NBs (n=23 cells from 5 brains) and polyploid, *sas4*^*mut*^ NBs (n=56 cells from 22 brains). (D) Dot plot showing the time spent in mitosis for Ctrl, *sas4*^*mut*^ NBs (n=23 cells from 5 brains) and polyploid, *sas4*^*mut*^ NBs (n=28 cells from 14 brains). (E) XY plot and corresponding linear regression between the number of nuclei generated at anaphase and polyploid, *sas4*^*mut*^ cell area (n=56 cells from 22 brains). (F) Stills of time-lapse movies of mitotic polyploid, *ncd*^*mut*^ NB expressing Tubulin-GFP (in green and grey in the bottom insets) and Histone 2B-RFP (in red). (G) Graph bars showing the percentage of cells presenting bipolar and multipolar mitosis in Ctrl, *ncd*^*mut*^ NB (n=23 cells from 3 brains) and polyploid, *ncd*^*mut*^ NBs (n=34 cells from 20 brains). (H) Dot plot showing the time spent in mitosis for Ctrl, *ncd*^*mut*^ NB (n=24 cells from 3 brains) and polyploid, *ncd*^*mut*^ NBs (n=15 cells from 12 brains). (I) XY plot and corresponding linear regression between the number of nuclei generated at anaphase and polyploid, *ncd*^*mut*^ cell area (n=34 cells from 20 brains). (A, B and F) Orange and white dotted circles surround cells and nuclei, respectively. Time of mitosis is represented in min.sec and time 00.00 corresponds to NEBD. Schematic representations of mitosis are shown above the stills. (D and H) Statistical significance was determined using a Mann-Whitney test. Lines represent the mean ± SD. (E and I) Statistical significance of the correlation was determined by a Spearman r test. R corresponding to the correlation coefficient. P corresponds to the p-value. Scale bar = 10µm.

We concluded that MT nucleation from the centrosomes greatly contributes to the assembly of multipolar spindles in polyploid NBs. However, in large NBs, where chromosome number is increased, mitotic spindles cannot resolve into a bipolar array even in the absence of centrosomes or with reduced centrosome numbers.

### Ncd/HSET is required, but is not sufficient to fully cluster spindle poles in polyploid NBs

Extra centrosomes can be generated in diploid NBs by Plk4/Sak over-expression (OE) [7]. In SakOE NBs as in other cells containing extra centrosomes, the minus-end directed kinesin Ncd (*Drosophila* ortholog of HSET) plays an essential role in promoting centrosome clustering [18, 31]. We tested whether Ncd was also contributing to the initial clustering observed in polyploid NBs. We induced polyploidy in *ncd* mutants (*ncd*^*mut*^) [32] and followed mitosis by time-lapse microscopy. We readily noticed that, after NEBD, in polyploid, *ncd*^*mut*^ NBs extra MT-nucleating sites did not cluster. Instead, they formed individual and independent poles, leading to the formation of a network of multiple mini-spindles all over the cell, which, interestingly, remained connected to each other (time 12.00 to 38.00 min and as depicted in schemes above, Figure 3F and Video 12). As cells progressed through mitosis, this multipolar status was maintained, and nuclear divisions were always multipolar, while mitosis was always bipolar in diploid, *ncd*^*mut*^ NBs (Figure 3G, Supplementary Figure 2D and Video 13). Although mitotic duration was increased in polyploid, *ncd*^*mut*^ NBs compared to diploid controls (47.36±23.16 min in polyploid, *ncd*^*mut*^ vs 10.32±1.23 min in *ncd*^*mut*^, p<0,0001), it was similar to polyploid NBs (Figure 3H and Supplementary Table 1), suggesting that failure in the initial step of centrosome clustering does not perturb MT nucleation or delay mitotic progression. The vast majority of polyploid, *ncd*^*mut*^ NBs presented a NI superior to 1 (Supplementary Figure 1F), confirming the increase in multipolarity.

Importantly, in polyploid, ncd^mut^ the regression slope between the number of nuclei generated at anaphase and cell area was quite high, even significantly higher than in polyploid NBs (Figure 3I, Supplementary Table 1, p<0,0001), suggesting that interfering with the initial centrosome clustering conditions the multipolarity outcome observed in polyploid, *ncd*^*mut*^.

To observe a link between the impact of MT nucleation pathways and genetic stability in polyploid cells, we quantified the percentage of NBs that generated micronuclei (MN) in each mutant background. MN are normally formed through chromosome missegregation and are prone to chromotripsis and genetic instability [33]. MN were identified based on their size as any nucleus smaller than the average diploid nucleus in Ctrl NBs. Importantly, while the combination of polyploidy with *ncd*^*mut*^ or Mars^KD^ resulted in an increase in the frequency of MN generated at anaphase, in polyploid, *sas4*^*mut*^ this frequency was reduced (Supplementary Table 1). We concluded that the absence of centrosomes in polyploid NBs represents an improvement not only for the assembly of bipolar spindles, but also for minimizing defects in chromosome attachment and segregation.

### Abnormal DNA content and shape inhibit spindle pole clustering

Since *ncd*^*mut*^ aggravated the multipolarity of polyploid mitosis, we hypothesized that Ncd could play an important role in setting up centrosome clustering during the initial stages of polyploid mitosis, thus promoting a reduction in the number of spindle poles (albeit insufficient to establish bipolarity). To test this hypothesis, we decided to characterize centrosomes in polyploid mitotic NBs. First, we performed a fixed analysis of late prometaphase NBs using centriole and PCM markers and second, we carried out time-lapse microscopy of brains expressing Histone-RFP and the centrosome component Spd-2 tagged with GFP (Spd2-GFP). We found that, in polyploid cells, several centrosomes could be clustered around different poles that surrounded highly condensed chromosomes (Figure 4A), which was never seen in diploid NBs with extra centrosomes [7]. Additionally, in time lapse movies, we observed dynamic initial clustering at NEBD that gathered centrosomes together in more than two poles, which remained isolated and separated from each other by the condensed chromosomes (Supplementary Figure 3A-B and Videos 14 and 15).

**Figure 4.**
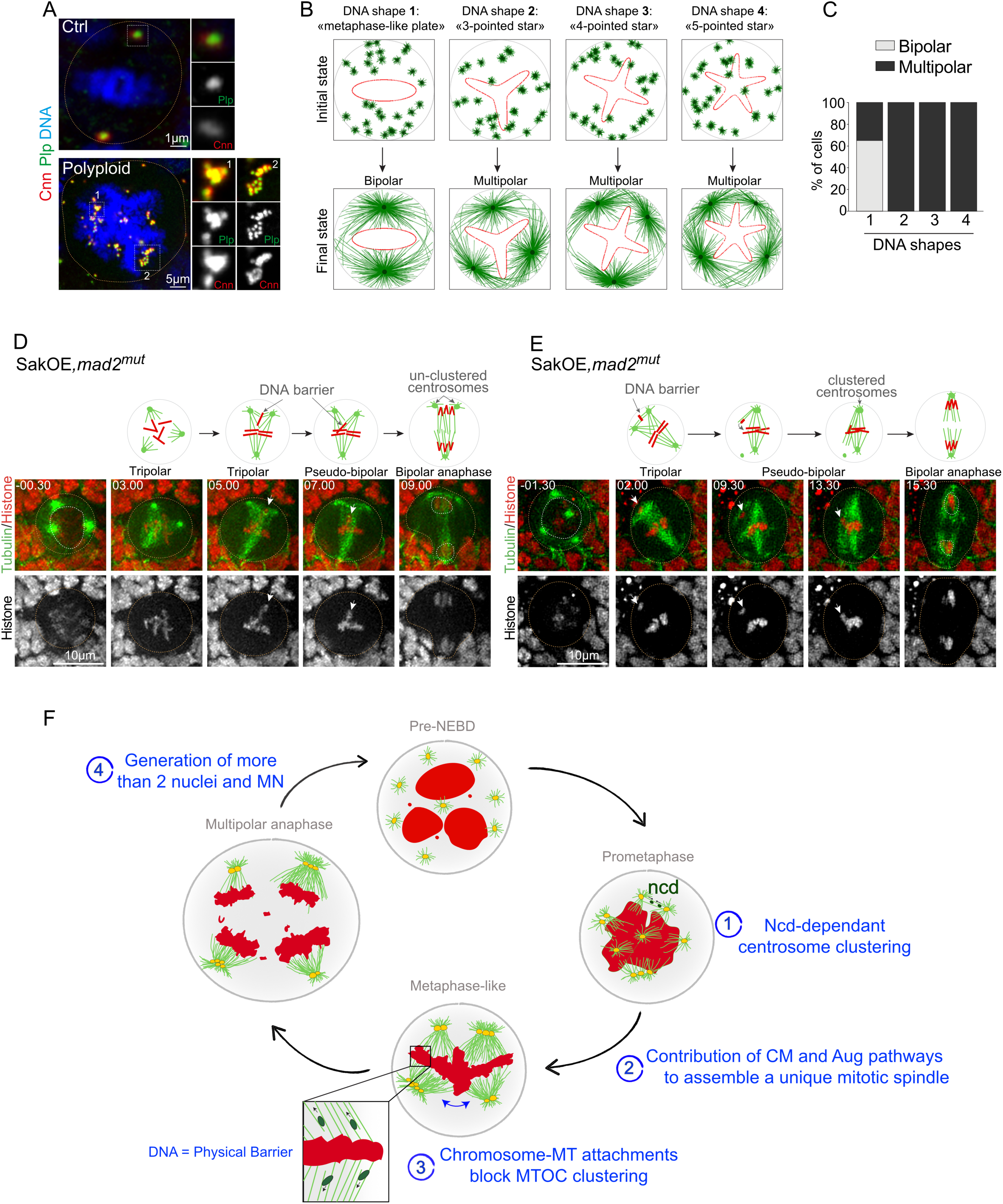
Extra-DNA acts as a physical barrier to spindle pole clustering. (A) Immunostaining images of Ctrl and polyploid NBs stained with a centriolar marker Plp (in green) and a PCM marker Cnn (red), also shown in grey in zoom insets from dotted white squares. The DNA is shown in blue and orange dotted circles surround cells. (B) Representative images of computational simulation, on the Cytosim software, of spindle formation in different DNA shape configurations. Cells present a fixed number of 30 MT-nucleating sites (centrosomes, shown in green) at initial state. The DNA (in red) shows different shape configurations (“bipolar metaphase-like plate”, “3-, 4- and 5-pointed star” shapes). As simulation proceeds, centrosomes cluster to form spindle poles. At final stage, only the DNA configuration “metaphase-like plate” presents a bipolar status. In contrast, more complex shapes assemble in a multipolar manner. (C) Graph bars showing the percentage of cells presenting bipolar and multipolar status in the four different DNA configuration shapes. (n=20 simulations per condition). (D-E) Stills of time-lapse movies of mitotic SakOE, *mad2*^*mut*^ NBs expressing Tubulin-GFP (in green) and Histone 2B-RFP (in red and grey in the bottom insets). Orange and white dotted circles surround cells and nuclei, respectively. Time of mitosis is indicated in min.sec and time 00.00 corresponds to NEBD. Schematic representations of mitosis are shown above the stills. (D) Diploid SakOE, *mad2*^*mut*^ NB with 3 MT-nucleating sites builds first a tripolar spindle. A DNA masse acts as a barrier blocking the clustering of two poles (white arrow). At anaphase onset, while centrosomes are still un-clustered, two nuclei are generated. (E) Diploid SakOE, *mad2*^*mut*^ NB presents several MT-nucleating sites and assembles a tripolar spindle containing a chromosome between two poles (white arrow). As mitosis proceeds, the chromosome is repositioned toward the metaphase plate and a pseudo-bipolar spindle forms. At bipolar anaphase, two nuclei are generated. (F) Polyploidy induced by cytokinesis failure generates large cells containing several nuclei and centrosomes. **1**. At NEBD, extra-centrosomes in close proximity to each other start to cluster together through a Ncd-dependant mechanism. **2**. The CMP and Augmin pathways contribute to the formation of a single multipolar spindle. **3**. The presence of extra-DNA mass inhibits the full clustering of the extra poles into two main poles, by acting as a physical barrier. **4**. At anaphase, several nuclei and micronuclei are generated.

So far, our results show that an increase in DNA and MT nucleation capacity is incompatible with the assembly of a bipolar mitotic spindle in polyploid cells. The presence of multiple centrosomes likely fuels an important fraction of the excessive MT nucleation. While initial centrosome clustering is achieved through the activity of Ncd, this activity is not sufficient to gather all centrosomes into two main spindle poles. Taken together, our results suggest that the increased DNA content of polyploid NBs might block the completion of spindle pole clustering, preventing the formation of a bipolar mitotic spindle. We tested whether DNA could function as a physical barrier that could maintain multiple spindle poles away from each other. Indeed, it has been shown that Ncd requires a minimal distance to promote clustering through cross-linking of antiparallel MTs [31]. We first established an in-silico approach. We designed simulations using Cyto-sim, a cytoskeleton-dedicated agent-based software [34]. We configured a minimalist system of spindle formation composed of a fixed centrosome number, microtubules, Ncd-like motors and DNA plates displaying different morphologies (based on DNA configurations seen in polyploid NBs, *in vivo*). While centrosomes clustered into two main poles when the DNA shape corresponded to a classical metaphase plate, simulations where the DNA shape was more complex generated, in most of the cases, a multipolar outcome (Figure 4B-C and Videos 16-19). These observations suggest that DNA shape impacts multipolarity, *in silico*. To strengthen our hypothesis, we then tried to decrease DNA content of polyploid NBs *in vivo* using DNAse treatment and laser ablation. Unfortunately, neither of these treatments was useful as they were highly toxic for the brain (data not shown). To overcome this problem, we designed an inverse strategy and searched for diploid conditions where DNA could inhibit centrosome clustering. While, extra-centrosomes in diploid SakOE NBs efficiently cluster to form a bipolar spindle, in combination with mutations in the spindle checkpoint *mad2* gene, a fraction of NBs divide multipolarly [18, 7, 35]. Careful analysis of mitotic SakOE, *mad2*^*mut*^ NBs by time-lapse microscopy revealed that in certain diploid cells, chromosomes can block centrosome clustering (Figure 4D-E - white arrows and as depicted in schemes above -and Movies 20-21). However, if the DNA is repositioned toward the metaphase plate, a bipolar spindle is assembled. (Figure 4E and Movie 20). In other NBs, centrosomes were maintained in an un-clustered status until anaphase (Figure 4D, Movie 21). These observations strengthen our hypothesis and support a model in which chromosomes act as a barrier to spindle pole clustering.

## Discussion

Here, we have investigated the contribution of different MT nucleation pathways to multipolar division of polyploid cells. We found that while the chromatin-mediated and Augmin pathways contribute to the establishment of a single mitotic spindle, even if multipolar, the MTs generated by these pathways do not seem to prevent the establishment of bipolarity. In contrast, centrosomes seem to be major contributors to spindle multipolarity, since centrosome removal rescued bipolar spindle assembly at least in small-medium sized polyploid NBs. Our data is consistent with a model where the initial centrosome clustering, occurring during prometaphase, is achieved through the activity of Ncd/Hset (Figure 4F). Importantly, in diploid NBs that contain extra centrosomes, Ncd is essential and sufficient to sustain bipolar spindle assembly [7]. Our results suggest that the accumulation of excessive DNA-which is then found in the form of condensed mitotic chromosomes-acts as a physical barrier to the maturation of complete spindle pole clustering, maintaining multiple MT-organizing centers (MTOCs) distant from each other. It has been shown that DNA shape and size can influence the morphology of the mitotic spindle assembled on chromatin coated beads [36]. Our work suggests that a similar phenomenon might be taking place in polyploid cells *in vivo* and that the presence of increased condensed DNA contributes to the maintenance of multipolarity.

Our work provides a framework to understand the generation of CIN in cells that have failed cytokinesis and, more importantly, how aneuploid karyotypes can be established after WGDs. Importantly, cytokinesis failure might also cause other type of pathologies. Mutations in MKLP1, the human Pav homolog have been identified in human populations suffering of congenial dysenrythropoetic anemia type III (CDA III), where erythroblasts show features such as multinucleation and nuclei size variation [37]. It will be important to understand whether tissue specific characteristics allow these cells to be maintained and how they contribute to anemia in this particular case.

## Acknowledgment

We acknowledge the PiCT-IBiSA platform and Nikon Imaging Center at Institut Curie for providing microscopes and V. Fraisier for outstanding help with image acquisition and advice. We thank V. Marthiens, D. Vargas-Hurtado, S. Gemble, D. Fachinetti, F. Edwards, O. Goundiam and S. Passemard for helpful discussion and/or comments on the manuscript. This work was supported by an ERC CoG (ChromoNumber-LS3, ERC-2016-COG), Institut Curie and CNRS. M.N was funded by La Ligue contre le cancer and FRM (FDT20160435352) grants and A.G by an FRM (ECO20170637529) fellowship. GL was funded by ANR (ANR-16-CE13 to M.-E. Terret).

## Author contribution

A.G performed most of the experiments, analyzed the data and generated the figures. M.N conceived the project, performed experiments and established the majority of *Drosophila* lines and tools. G.L generated the computational models and contributed to the discussions. C.P generated certain tools and helped with fly pushing. R.B conceived the project, wrote the manuscript and supervised the project. A.G., M.N. and R.B. interpreted the data which was discussed between all authors during the preparation of the manuscript.

## Methods

### CONTACT FOR REAGENT AND RESOURCE SHARING

Further information and requests for resources and reagents should be directed to and will be fulfilled by the Lead Contact, Renata Basto

### EXPERIMENTAL MODEL AND SUBJECT DETAILS

Flies were raised in plastic vials or bottles containing homemade *Drosophila* culture medium (0,75% agar, 3,5% organic wheat flour, 5% yeast, 5,5% sugar, 2,5% nipagin, 1% penicillin-streptomycin and 0,4% propanic acid). Fly stocks were conserved at 18°C. For temporal activation of RNAi, crosses were kept at 18°C and switched to 29°C for 24 to 48 hours. Other experimental crosses were maintained at 25°C.

### METHOD DETAILS

#### Live-imaging

Mid third-instar larval (L3) brains were dissected in Schneider’s *Drosophila* medium supplemented with 10% heat-inactivated fetal bovine serum, Penicillin (100 units.ml-1) and Streptomycin (100 µg.ml-1). Several brains were placed on a glass bottom dish with near 10 µl of medium, covered with a permeable membrane and sealed around the membrane borders with oil 10 S Voltalef.

#### Immunofluorescence on squashed brains

L3 brains were dissected in PBS and fixed for 20 to 40 min at room temperature (RT) in 4% formaldehyde diluted in PBS. After fixation, brains were transferred in a drop of 45% acetic acid on a coverslip for 15 seconds and then immediately moved in a drop of 60% acetic acid. Acetic acid was diluted in water. A slide was then placed on top of the coverslip and brains were squashed with a pencil until the tissue appeared transparent. The slide was rapidly flash-frozen in liquid nitrogen before the removal of the coverslip using a sharp blade. Slides were incubated 7 minutes in −20°C methanol, washed and permeabilized 3 times for 10 min in PBST 0,1% solution (PBS, 0,1% Triton X-100 and 0,02% Sodium Azide) at RT. Slides were dried at RT before incubation with 10 µL of primary antibodies diluted in PBT 0,1% and covered with a coverslip in a humid chamber at 4°C overnight (O/N). After removal of the coverslips, slides were washed 3 times for 10 min in PBST 0,1% at RT. 10 µL of secondary antibody dilution was added to the slides; slides were covered with coverslips in a dark, humid chamber and incubated 3h at 25°C. Slides were washed 3 times for 10min in PBST 0,1% and incubated in 0.5µg.ml^-1^ Hoechst for 15min at RT. Slides were mounted using 22×22 mm coverslip using 10 µL of mounting medium (1.25%n-Propyl Gallate, 75% glycerol, 25% H20).

#### Immunofluorescence on whole mount brains

Mid third-instar larval brains were dissected in PBS and fixed for 30 min at RT in 4% paraformaldehyde diluted in PBS. After fixation, brains were washed and permeabilized 3 times for 10 min in PBST 0,3% (PBS, 0,3% Triton X-100 and 0,02% Sodium Azide). Brains were then incubated in primary antibodies diluted in PBST 0,3%, at 4°C O/N in a humid chamber. After 3 times 10min washes in PBST 0,3%, brains were incubated in secondary antibodies and DAPI dilution (in PBST 0,3%), either for several hours at 25°C or O/N at 4°C, protected from light into a humid chamber. After 3 washes for 10min in PBST 0,3%, brains were rinsed in PBS and mounted using 12 mm coverslips with 5 µL of mounting medium (1.25%n-Propyl Gallate, 75% glycerol, 25% H20).

#### Image acquisition and processing

For live-imaging, images were acquired with 60X oil objective (NA 1.4) on two microscopes: an Inverted Spinning Disk Confocal Roper/Nikon (a Yokagawa CSU-X1 spinning head mounted on a Nikon Ti-E inverted microscope equipped with a camera EMCCD 512×512 Evolve (Photometrics)) and the wide-field Inverted Spinning Disk Confocal Gattaca/Nikon (a Yokagawa CSU-W1 spinning head mounted on a Nikon Ti-E inverted microscope equipped with a camera sCMOS (complementary metal-oxide semiconductor) 1200×1200 Prime95B (Photometrics)). For both microscopes, images were acquired at time intervals spanning from 30sec (diploid conditions) to 2min (polyploid conditions) and 30 to 50 Z-stacks of 1 to 1.5 µm controlled by the Metamorph software.

For squashed brain experiments, images were acquired with 40X (NA 1.25), 63X (NA 1.32) or 100X (NA 1.4) oil objectives on an Epifluorescent Upright microscope Leica DM6000 equipped with a camera CCD, 1392×1040 CoolSnap HQ (Photometrics). Intervals for Z-stacks acquisitions were set to 0.3 to 1µm using the Metamorph software.

For whole mount brain experiments, images were acquired on a confocal Nikon A1R Ti-E inverted microscope with a 60X (NA 1.4) oil objective. Interval for Z-stack acquisitions was set up with 0.3µm using the NIS Element software.

#### Computational simulations

Simulations were performed using Cytosim (www.cytosim.org)[38].

As cells were mostly spherical during mitosis, we confined the MTs and motors in a circular shape (2D). Centrosomes were modelled as asters nucleating MTs, radially. Initially, centrosomes were randomly placed inside the cell and outside the chromatin plate, corresponding to the experimental situations just after NEBD. MTs were considered as flexible polymers with static minus-ends and dynamic plus-ends that could grow and shrink according to the dynamic instability model. The kinesin motors Ncd were represented as an entity composed of 2 separated binding MT « heads », one static non-motor domain and one motor-domain that can move towards the minus-end of the MT [39]. Ncd was shown to bind and slide to anti-parallel MTs and crosslink the parallel ones [40]. Here, as we were only interested in the clustering effect (thus the antiparallel sliding), we constrained Ncd-like entities to only bind to antiparallel MTs that reduced the amount of entities to simulate. Finally, we simulated the metaphase plate as a static obstacle (an area of a defined shape) that MTs could not cross. To simplify the model, we did not consider MT nucleation from the plate neither deformation of the plate by MTs. MTs were excluded from this area and were prevented from sliding on the surface by the presence of plus-end tracking entities (mimicking chromokinesins present on chromosomes or capture by prometaphase kinetochores) similar to simulations from [41].

Parameters in the simulations were chosen to match the experimental conditions when measurable or from the literature when available (Supplemental Table 2). Note that we varied the values of these parameters to check that our numerical conclusions were not greatly affected by this choice.

### QUANTIFICATION AND STATISTICAL ANALYSIS

Image analysis and quantifications were performed using Fiji software. To compare the same range of polyploid cells for all conditions, quantifications were done on cells with an area inferior to 6000µm2. Centrosomes were manually counted and identified as a colocalization of Cnn and Plp markers on squashed brains experiments. The bipolar/multipolar state of the mitotic spindle, the number of nuclei, the cell area and the mitotic timing were assessed on live-imaging data. To calculate the nuclear index (number of nuclei at anaphase vs mitotic entry), the number of nuclei and micronuclei were manually counted for all Z-stacks at pre-nuclear envelop breakdown (NEBD) and at anaphase, when nuclei are individualized and before chromosomes decondense (see white dotted circles on Figures). A nucleus was considered as micronucleus when the diameter was smaller than an average diploid nucleus (calculated as the mean diameter of 5 WT cell nuclei, independently on both microscopes, 31,8pixels and 40,4px at pre-NEBD and 15,6px and 19,4px at anaphase, for the Spinning Disk Confocal Roper and Gattaca, respectively). Knowing the time intervals between acquired frames, the mitotic timing was calculated from NEBD to anaphase onset. The cell area was calculated using the “Freehand selection” tool on Fiji software to draw cell contour. All statistical analyses were performed on Prism software (See Figure legends for details). For the analysis of the correlation between the number of nuclei at anaphase and NB area, the correlation coefficient *r* corresponds to the strength of the relationship between both variables and the regression slope represents the rate of change, thus to which extend the number of nuclei increases with cell area. For figures, images were processed on Photoshop and Affinity Photo, and mounted using Illustrator and Affinity Designer.

### KEY RESOURCES TABLE

| REAGENT or RESOURCE | SOURCE | IDENTIFIER |
| --- | --- | --- |
| **Antibodies** |  |  |
| Guinea Pig anti-Cnn (1:1000) | Raff J.W. Lab [42] | N/A |
| Rabbit anti-D-PLP (PPHA + PPHC) (1:1000 each) | Raff J.W. Lab [43] | N/A |
| Mouse anti-α-Tubulin (DM1α) (1 :500) | Sigma-Aldrich | Cat# T6199<br>RRID:AB_477583 |
| Goat anti-Rabbit IgG Alexa Fluor 488 (1:250) | Thermo Fisher Scientific | Cat# A-11034,<br>RRID:AB_2576217 |
| Goat anti-Guinea Pig IgG Alexa Fluor 647 (1:250) | Thermo Fisher Scientific | Cat# A-21450,<br>RRID:AB_2735091 |
| Goat anti-Mouse IgG Alexa Fluor 568 (1:250) | Thermo Fisher Scientific | Cat# A-11031,<br>RRID:AB_144696 |
| **Chemicals, Peptides, and Recombinant Proteins** |  |  |
| 4',6-diamidino-2-phenylindole DAPI | Thermo Fisher Scientific | Cat# 62248 |
| Hoechst | Thermo Fisher Scientific | Cat# 33342 |
| Phosphate-Buffered Saline (PBS) | VWR | Cat# L182-10 |
| 16% Paraformaldehyde EM Grade | EMS | Cat# 15710 |
| 37% Formaldehyde, Microfiltered | EMS | Cat# 15686 |
| Acetic acid 99-100% | VWR Chemicals | Cat# 20103.295 |
| Methanol RPE | CARLO ERBA Reagents | Cat# 414819 |
| Triton X-100 | Euromedex | Cat# 2000-C |
| Sodium Azide | Sigma-Aldrich | Cat# 71289 |
| Schneider's *Drosophila* Medium | Gibco | Cat# 21720024 |
| Foetal Bovine Serum, heat inactivated | Gibco | Cat# 10500 |
| Penicillin-Streptomycin | Gibco | Cat# 15140 |
| 35 mm glass bottom Dish | MatTek Corporation | Cat# P35G-1.5-14-C |
| Membrane kit, Standard | YSI | Cat# 098094 |
| Oil 10 S, VOLTALEF® | VWR Chemicals | N/A |
| **Experimental Models: Organisms/Strains** |  |  |
| w[118] | BDSC | RRID:BDSC_5905 |
| WT isogenic | Basto Lab | N/A |
| Actin-GAL4 (act5C-GAL4) | Bloomington Drosophila Stock Center | RRID:BDSC_3954 |
| Asense-GAL4 | Lee Lab [44] | N/A |
| GAL80<sup>ts</sup> (α-Tub84B-GAL80<sup>ts</sup>) | BDSC | RRID:BDSC_7017 |
| α-Tubulin-GFP | Raff J.W Lab | N/A |
| Histone-RFP (H2Av-mRFP) | BDSC | RRID:BDSC_23651 |
| *sqh<sup>l</sup>* / FM7, Kruppel-GAL4,UAS-GFP | Karess R.E Lab | N/A |
| UAS-pav<sup>RNAi</sup> | BDSC | RRID:BDSC_42573 |
| UAS-dgt2<sup>RNAi</sup> | BDSC | RRID:BDSC_31729 |
| UAS-mars<sup>RNAi</sup> | BDSC | RRID:BDSC_33929 |
| *DSas4<sup>mut(S2214)</sup>* | BDSC | RRID:BDSC_12119 |
| *ncd<sup>mut</sup> (ncd<sup>l</sup>)* | [32] | N/A |
| pUbqSak-#68 (SakOE) | [7] | N/A |
| *mad2<sup>mut</sup>* | BDSC | 22495 |
| Asense-GAL4,Tub-GFP | Basto Lab | N/A |
| UAS-Pav<sup>RNAi</sup>,Hist-RFP | Basto Lab | N/A |
| Tubulin-GFP,Histone-RFP | Basto Lab | N/A |
| Actin-GAL4,GAL80<sup>ts</sup> | Bellaiche Lab | N/A |
| Tub-GFP,Hist-RFP ; UAS-dgt2<sup>RNAi</sup> | This study | N/A |
| Tub-GFP,Hist-RFP ; UAS-mars<sup>RNAi</sup> | This study | N/A |
| Asense-GAL4,Tub-GFP; *sas4<sup>mut</sup>* | This study | N/A |
| UAS-Pav<sup>RNAi</sup>,Hist-RFP ; *sas4<sup>mut</sup>* | This study | N/A |
| Asense-GAL4,Tub-GFP; *ncd<sup>mut</sup>* | This study | N/A |
| UAS-Pav<sup>RNAi</sup>,Hist-RFP ; *ncd<sup>mut</sup>* | This study | N/A |
| *sqh<sup>l</sup>* ; Tub-GFP,Hist-RFP | This study | N/A |
| Tub-GFP, Hist-RFP; *mad2<sup>mut</sup>* | [35] | N/A |
| pUbqSak-#68, *mad2<sup>mut</sup>* | [35] | N/A |
| **Software and Algorithms** |  |  |
| Fiji | Open Sources | https://fiji.sc/ |
| Prism 7 | GraphPad software, Inc. | Version 7.0a |
| Photoshop | Adobe | Version 2017.0.1 |
| Affinity Photo and Designer | Serif | N/A |
| Illustrator | Adobe | Version 2017.0.2 |
| Cytosim | [34] | www.cytosim.org |
| Metamorph | Molecular Devices, USA | N/A |
| NIS Element software | Nikon | N/A |

| Other |  |  |
| --- | --- | --- |
| Cover glasses Menzel gläser, 18x18 mm | VWR | Cat# 631-1331 |
| Cover glasses, square, 22x22 mm | VWR | Cat# 631-0125 |
| Cover glasses, circular, 12 mm Ø | Marienfeld Superior | Cat# 0111520 |
| Slide Superfrost cut edges | Thermo Fisher Scientific | Cat#<br>AA00008232E00MNT10 |

## References

1. Zack, T.I., Schumacher, S.E., Carter, S.L., Cherniack, A.D., Saksena, G., Tabak, B., Lawrence, M.S., Zhsng, C.Z., Wala, J., Mermel, C.H., et al. (2013). Pan-cancer patterns of somatic copy number alteration. Nat Genet 45, 1134–1140.

2. Bielski, C.M., Zehir, A., Penson, A.V., Donoghue, M.T.A., Chatila, W., Armenia, J., Chang, M.T., Schram, A.M., Jonsson, P., Bandlamudi, C., et al. (2018). Genome doubling shapes the evolution and prognosis of advanced cancers. Nat Genet 50, 1189– 1195.

3. Fujiwara, T., Bandi, M., Nitta, M., Ivanova, E.V., Bronson, R.T., and Pellman, D. (2005). Cytokinesis failure generating tetraploids promotes tumorigenesis in p53-null cells. Nature 437, 1043–1047.

4. Storchova, Z., and Pellman, D. (2004). From polyploidy to aneuploidy, genome instability and cancer. Nat Rev Mol Cell Biol 5, 45–54.

5. Marthiens, V., Piel, M., and Basto, R. (2012). Never tear us apart--the importance of centrosome clustering. J Cell Sci 125, 3281–3292.

6. Boveri, T. (2008). Concerning the origin of malignant tumours by Theodor Boveri. Translated and annotated by Henry Harris. J Cell Sci 121 Suppl 1, 1–84.

7. Basto, R., Brunk, K., Vinadogrova, T., Peel, N., Franz, A., Khodjakov, A., and Raff, J.W. (2008). Centrosome amplification can initiate tumorigenesis in flies. Cell 133, 1032–1042.

8. Sercin, O., Larsimont, J.C., Karambelas, A.E., Marthiens, V., Moers, V., Boeckx, B., Le Mercier, M., Lambrechts, D., Basto, R., and Blanpain, C. (2016). Transient PLK4 overexpression accelerates tumorigenesis in p53-deficient epidermis. Nature cell biology 18, 100–110.

9. Levine, M.S., Bakker, B., Boeckx, B., Moyett, J., Lu, J., Vitre, B., Spierings, D.C., Lansdorp, P.M., Cleveland, D.W., Lambrechts, D., et al. (2017). Centrosome Amplification Is Sufficient to Promote Spontaneous Tumorigenesis in Mammals. Developmental cell 40, 313–322 e315.

10. Fox, D.T., Gall, J.G., and Spradling, A.C. (2010). Error-prone polyploid mitosis during normal Drosophila development. Genes Dev 24, 2294–2302.

11. Duncan, A.W., Taylor, M.H., Hickey, R.D., Hanlon Newell, A.E., Lenzi, M.L., Olson, S.B., Finegold, M.J., and Grompe, M. (2010). The ploidy conveyor of mature hepatocytes as a source of genetic variation. Nature 467, 707–710.

12. Dewhurst, S.M., McGranahan, N., Burrell, R.A., Rowan, A.J., Gronroos, E., Endesfelder, D., Joshi, T., Mouradov, D., Gibbs, P., Ward, R.L., et al. (2014). Tolerance of whole-genome doubling propagates chromosomal instability and accelerates cancer genome evolution. Cancer Discov 4, 175–185.

13. Nano, M., Simon, A., Gogendeau, D., Pennetier, C., Fraisier, V., Marthiens, V., and Basto, R. Cell cycle asynchrony generates DNA damage at mitotic entry in polyploid cells.

14. Karess, R.E., Chang, X.J., Edwards, K.A., Kulkarni, S., Aguilera, I., and Kiehart, D.P. (1991). The regulatory light chain of nonmuscle myosin is encoded by spaghetti-squash, a gene required for cytokinesis in Drosophila. Cell 65, 1177–1189.

15. Adams, R.R., Tavares, A.A., Salzberg, A., Bellen, H.J., and Glover, D.M. (1998). pavarotti encodes a kinesin-like protein required to organize the central spindle and contractile ring for cytokinesis. Genes Dev 12, 1483–1494.

16. Ikeshima-Kataoka, H., Skeath, J.B., Nabeshima, Y., Doe, C.Q., and Matsuzaki, F. (1997). Miranda directs Prospero to a daughter cell during Drosophila asymmetric divisions. Nature 390, 625–629.

17. Homem, C.C., and Knoblich, J.A. (2012). Drosophila neuroblasts: a model for stem cell biology. Development 139, 4297–4310.

18. Kwon, M., Godinho, S.A., Chandhok, N.S., Ganem, N.J., Azioune, A., Thery, M., and Pellman, D. (2008). Mechanisms to suppress multipolar divisions in cancer cells with extra centrosomes. Genes Dev 22, 2189–2203.

19. Musacchio, A. (2015). The Molecular Biology of Spindle Assembly Checkpoint Signaling Dynamics. Current biology: CB 25, R1002–1018.

20. Hayward, D., Metz, J., Pellacani, C., and Wakefield, J.G. (2014). Synergy between multiple microtubule-generating pathways confers robustness to centrosome-driven mitotic spindle formation. Developmental cell 28, 81–93.

21. Prosser, S.L., and Pelletier, L. (2017). Mitotic spindle assembly in animal cells: a fine balancing act. Nat Rev Mol Cell Biol 18, 187–201.

22. Goshima, G., Wollman, R., Goodwin, S.S., Zhang, N., Scholey, J.M., Vale, R.D., and Stuurman, N. (2007). Genes required for mitotic spindle assembly in Drosophila S2 cells. Science 316, 417–421.

23. Goshima, G., Mayer, M., Zhang, N., Stuurman, N., and Vale, R.D. (2008). Augmin: a protein complex required for centrosome-independent microtubule generation within the spindle. The Journal of cell biology 181, 421–429.

24. Sanchez-Huertas, C., and Luders, J. (2015). The augmin connection in the geometry of microtubule networks. Current biology: CB 25, R294–299.

25. Yang, C.P., and Fan, S.S. (2008). Drosophila mars is required for organizing kinetochore microtubules during mitosis. Exp Cell Res 314, 3209–3220.

26. Kellogg, D.R., Moritz, M., and Alberts, B.M. (1994). The centrosome and cellular organization. Annu Rev Biochem 63, 639–674.

27. Conduit, P.T., Wainman, A., and Raff, J.W. (2015). Centrosome function and assembly in animal cells. Nat Rev Mol Cell Biol 16, 611–624.

28. Basto, R., Lau, J., Vinogradova, T., Gardiol, A., Woods, C.G., Khodjakov, A., and Raff, J.W. (2006). Flies without centrioles. Cell 125, 1375–1386.

29. Bettencourt-Dias, M., Rodrigues-Martins, A., Carpenter, L., Riparbelli, M., Lehmann, L., Gatt, M.K., Carmo, N., Balloux, F., Callaini, G., and Glover, D.M. (2005). SAK/PLK4 is required for centriole duplication and flagella development. Current biology: CB 15, 2199–2207.

30. Kirkham, M., Muller-Reichert, T., Oegema, K., Grill, S., and Hyman, A.A. (2003). SAS-4 is a C. elegans centriolar protein that controls centrosome size. Cell 112, 575– 587.

31. Rhys, A.D., Monteiro, P., Smith, C., Vaghela, M., Arnandis, T., Kato, T., Leitinger, B., Sahai, E., McAinsh, A., Charras, G., et al. (2018). Loss of E-cadherin provides tolerance to centrosome amplification in epithelial cancer cells. The Journal of cell biology 217, 195–209.

32. Endow, S.A., and Komma, D.J. (1998). Assembly and dynamics of an anastral:astral spindle: the meiosis II spindle of Drosophila oocytes. J Cell Sci 111 (Pt 17), 2487–2495.

33. Hintzsche, H., Hemmann, U., Poth, A., Utesch, D., Lott, J., and Stopper, H. (2017). Fate of micronuclei and micronucleated cells. Mutat Res 771, 85–98.

34. Francois, N., and Dietrich, F. (2007). Collective Langevin dynamics of flexible cytoskeletal fibers. New Journal of Physics 9, 427.

35. Gogendeau, D., Siudeja, K., Gambarotto, D., Pennetier, C., Bardin, A.J., and Basto, R. (2015). Aneuploidy causes premature differentiation of neural and intestinal stem cells. Nature communications 6, 8894.

36. Dinarina, A., Pugieux, C., Corral, M.M., Loose, M., Spatz, J., Karsenti, E., and Nedelec, F. (2009). Chromatin shapes the mitotic spindle. Cell 138, 502–513.

37. Liljeholm, M., Irvine, A.F., Vikberg, A.L., Norberg, A., Month, S., Sandstrom, H., Wahlin, A., Mishima, M., and Golovleva, I. (2013). Congenital dyserythropoietic anemia type III (CDA III) is caused by a mutation in kinesin family member, KIF23. Blood 121, 4791–4799.

38. Nedelec, F., and Foethke, D. (2007). Collective Langevin dynamics of flexible cytoskeletal fibers. New Journal of Physics 9, 427–427.

39. Letort, G., Bennabi, I., Dmitrieff, S., Nedelec, F., Verlhac, M.H., and Terret, M.E. (2019). A computational model of the early stages of acentriolar meiotic spindle assembly. Molecular biology of the cell, mbcE18100644.

40. Fink, G., Hajdo, L., Skowronek, K.J., Reuther, C., Kasprzak, A.A., and Diez, S. (2009). The mitotic kinesin-14 Ncd drives directional microtubule-microtubule sliding. Nature cell biology 11, 717–723.

41. Lacroix, B., Letort, G., Pitayu, L., Salle, J., Stefanutti, M., Maton, G., Ladouceur, A.M., Canman, J.C., Maddox, P.S., Maddox, A.S., et al. (2018). Microtubule Dynamics Scale with Cell Size to Set Spindle Length and Assembly Timing. Developmental cell 45, 496–511.e496.

42. Lucas, E.P., and Raff, J.W. (2007). Maintaining the proper connection between the centrioles and the pericentriolar matrix requires Drosophila centrosomin. The Journal of cell biology 178, 725–732.

43. Martinez-Campos, M., Basto, R., Baker, J., Kernan, M., and Raff, J.W. (2004). The Drosophila pericentrin-like protein is essential for cilia/flagella function, but appears to be dispensable for mitosis. The Journal of cell biology 165, 673–683.

44. Zhu, S., Lin, S., Kao, C.F., Awasaki, T., Chiang, A.S., and Lee, T. (2006). Gradients of the Drosophila Chinmo BTB-zinc finger protein govern neuronal temporal identity. Cell 127, 409–422.

