## Supplementary Information for "Chromosomes function as a barrier to mitotic spindle bipolarity in polyploid cells"

### Goupil, Nano et al. Supplementary Information

#### Supplementary figure legends

**Supplementary Figure 1: Polyploid brains present NBs with different level of polyploidization.**

(A) Immunostaining images of whole mount diploid and polyploid brain lobes stained with antibodies against $\alpha$-tubulin (in red) and Cnn (in green), to label mitotic spindle and pericentriolar material, respectively. DNA is shown in blue. Mitotic diploid and polyploid NBs are indicated in zoom insets. Different degrees of polyploidy are present in polyploid brain, highlighted by the cell size, centrosome number and DNA content. (B) Dot plot showing the number of centrosomes per cell in diploid Ctrl (n=30 NBs from 4 brain lobes) and polyploid ($Pav^{KD}$) (n=38 NBs from 10 brain lobes). Statistical significance was determined using a t-test. Lines represent the mean ± SD. (C) Stills of time-lapse movies of a mitotic polyploid NB expressing Tubulin-GFP (in green and grey in the bottom insets) and Histone 2B-RFP (in red). Orange and white dotted circles surround cells and nuclei, respectively. Time of mitosis is represented in min.sec and time 00.00 corresponds to NEBD. Low polyploidy level $Pav^{KD}$ NB, while superior to diploid NB, shows small size and low number of MT-nucleating sites. (D) XY plot representing the mitotic timing (min) related to polyploid cell area (n=60 cells from 31 brains). Statistical not significance (ns) of the correlation was determined by a Spearman r test. R corresponding to the correlation coefficient. (E) Schematic representation of the NI calculated as the ratio between the number of nuclei at anaphase versus at mitotic entry (pre-NEBD). Cells and nuclei are represented in green and red, respectively. In mitotic diploid cell, a unique nucleus divides in two daughter nuclei, the NI is then equal to 2. A polyploid cell with a NI inferior to 1 presents a number of nuclei at anaphase (e.g. 3) that is less than at mitotic entry (e.g. 4), while a polypoid cell with a NI between 1 and 2 or superior to 2 ends mitosis with more nuclei (e.g. 7) or more than the double (e.g. 10) than at mitotic entry (e.g. 4). (F) Graph bars showing the percentage of cells presenting a NI inferior or equal to 1 (Category (a): $NI \leq 1$), between 1 and 2 (Cat (b): $1< NI< 2$) or superior or equal to 2 (Cat (c): $NI \geq 2$) in diploid, Ctrl (n=34 cells from 2 brains), diploid, $Aug^{KD}$ (n=30 cells from 3 brains), diploid, $Mars^{KD}$ (n=13 cells from 1 brains), diploid, *sas4*$^{mut}$ (n=23 cells from 5 brains), diploid, *ncd*$^{mut}$ (n=23 cells from 3 brains), polyploid (n=54 cells from 28 brains), polyploid, $Aug^{KD}$ (n=24 cells from 10 brains), polyploid, $Mars^{KD}$ (n=44 cells from 26 brains), polyploid, *sas4*$^{mut}$ (n=27 cells from 14 brains) and polyploid, *ncd*$^{mut}$ (n=14 cells from 11 brains) NBs. Statistical significance was determined using a qhi$^2$ test for all conditions related to the corresponding ploidy control and between diploid and the corresponding polyploid conditions. Only significant values are shown. P corresponds to the p-value.

**Supplementary Table 1:**

Table showing the different parameters of mitotic NBs in diploid and polyploid conditions characterized here. The mitotic timing corresponds to the time elapsed from NEBD to anaphase onset in minutes (min). The nuclear index (NI) is calculated as the ratio between the number of nuclei at anaphase over mitotic entry. The slope of the linear regression between the number of nuclei generated at anaphase and polyploid cell area is determined using an equation of the linear regression line. The percentage (%) of cells with micronuclei (MN) corresponds to the percentage of cells generating at anaphase, at least one nucleus smaller in size than a diploid nucleus from Ctrl NBs.

Values correspond to the mean ± SD or percentage and the statistical significances are related to the ploidy control NBs (diploid Ctrl or polyploid). Statistical significances have been determined by Mann-Whitney test for mitotic timing and NI, by t-test for micronuclei and by slope comparison test of linear regression for the slope lines. P corresponds to the p-value, ns to "not significant" and ND to "not determined".

**Supplementary Figure 2: Disruption of MT-nucleating pathways and centrosome clustering factor Ncd do not perturb the bipolarity of the diploid mitotic spindle.**

(A-D) Stills of time-lapse movies of mitotic NBs expressing Tubulin-GFP (in green and grey in the bottom insets) and Histone 2B-RFP (in red). Orange and white dotted circles surround cells and nuclei, respectively. Time of mitosis is represented in min.sec and time 00.00 corresponds to NEBD. Schematic representations of mitosis are shown above the stills. (A) Diploid, Aug$^{KD}$ NB builts a normal bipolar spindle generating 2 nuclei at anaphase. (B) Diploid, Mars$^{KD}$ NB assembles a thinner bipolar spindle giving rise to two daughter cells after division. (C) In absence of centrosomes, diploid, *sas4*$^{mut}$ NB nucleates MTs from chromatin that elongates in an outward manner to form a bipolar spindle. Bipolar division gives rise to two daughter cells. (D) Diploid, *ncd*$^{mut}$ NB enter in mitosis with 2 centrosomes that assemble a bipolar spindle. At anaphase, the spindle bends but generates 2 nuclei. Scale bar=10µm.

**Supplementary Figure 3: In polyploid NBs, extra-centrosomes initially cluster together to form poles of the multipolar spindle.**

(A-B) Stills of time-lapse movies of mitotic NBs expressing Histone 2B-RFP (in red) and Spd2-GFP (in green and grey in the bottom insets), corresponding to DNA and centrosomes, respectively. Orange dotted circles surround cells. Time of mitosis is represented in min.sec and time 00.00 corresponds to NEBD. Schematic representations of mitosis are shown above the stills. Scale bar=10µm.

### Supplementary Figure 1

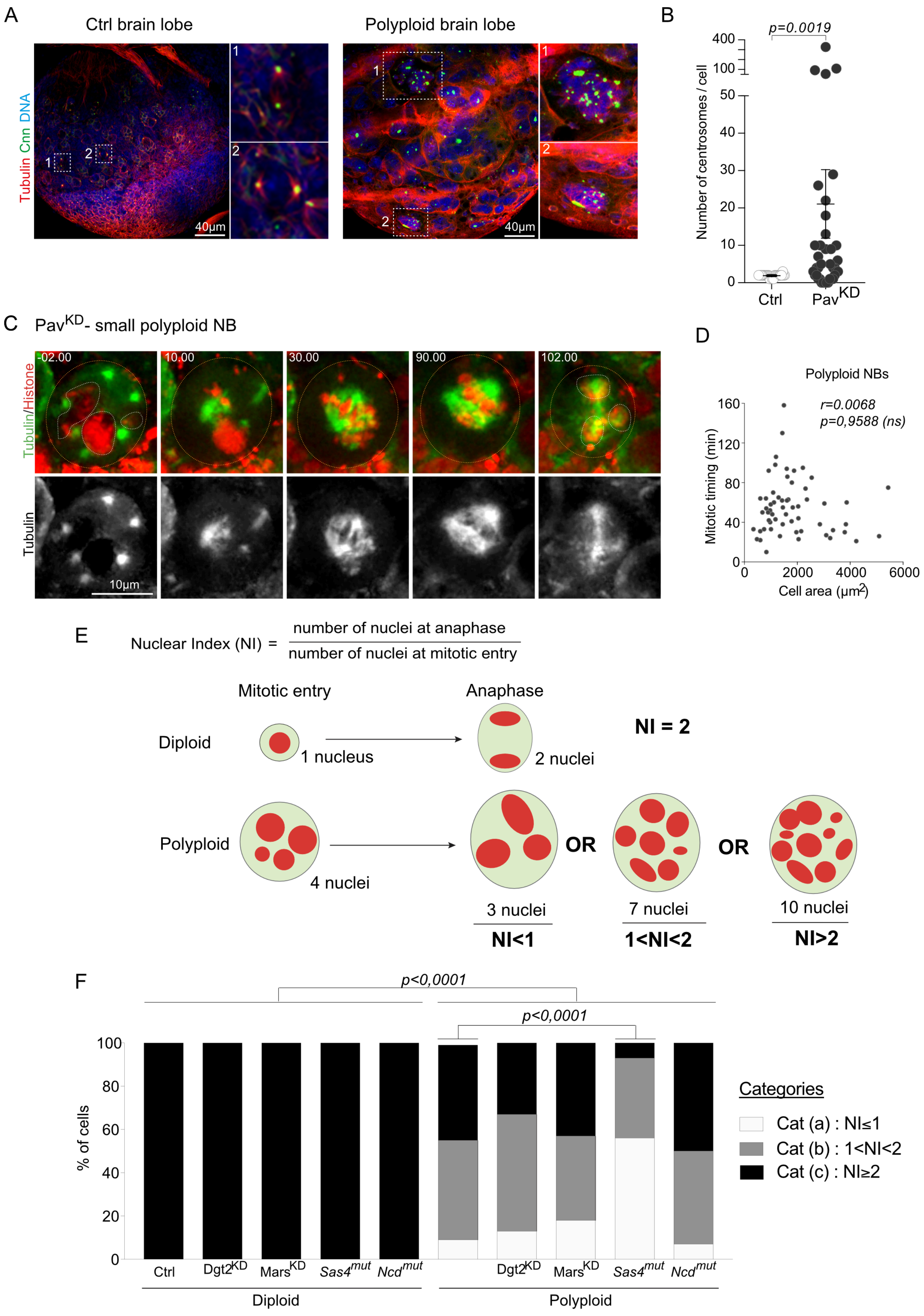

### Supplementary Table 1

| | Genotypes | Mitotic duration | | Nuclear Index (NI) | | Slope of linear regression | | Micronuclei (MN) | |
|---|---|---|---|---|---|---|---|---|---|
| | | Mean ± SD | *p-value* *Mann-Whitney test* | Mean ± SD | *p-value* *Mann-Whitney test* | Mean ± SD | *p-value* *slope comparison test* | % of cells | *p-value* *T-test* |
| Diploid | Diploid (Ctrl) | 07.37 ± 0.53 min | – | 2 ± 0 | – | – | – | 0% | – |
| | Diploid, Aug$^{KD}$ | 07.38 ± 3.07 min | *p<0.0001* | 2 ± 0 | *ND* | – | – | 0% | *ND* |
| | Diploid, Mars$^{KD}$ | 12.23 ± 4.04 min | *p<0.0001* | 2 ± 0 | *ND* | – | – | 0% | *ND* |
| | Diploid, *sas4*$^{mut}$ | 16.47 ± 8.34 min | *p<0.0001* | 2 ± 0 | *ND* | – | – | 0% | *ND* |
| | Diploid, *ncd*$^{mut}$ | 10.32 ± 1.23 min | *p<0.0001* | 2 ± 0 | *ND* | – | – | 0% | *ND* |
| Polyploid | Polyploid | 55.12 ± 28.07 min | – | 1.86 ± 0.64 | – | 0.0052 ± 0.0004 | – | 63% | – |
| | Polyploid, Aug$^{KD}$ | 71.30 ± 24.08 min | *p=0.0075* | 1.91 ± 1.23 | *p=0,3798(ns)* | 0.0060 ± 0.0008 | *p=0,2842(ns)* | 71% | *p =0,2994(ns)* |
| | Polyploid, Mars$^{KD}$ | 30.34 ± 12.32 min | *p<0.0001* | 1.76 ± 0.65 | *p=0,4694(ns)* | 0.0028 ± 0.0006 | *p=0.0002* | 90% | *p=0.0008* |
| | Polyploid, *sas4*$^{mut}$ | 38.19 ± 14.41 min | *p=0.0073* | 1.24 ± 0.42 | *p<0.0001* | 0.0021 ± 0.0003 | *p<0.0001* | 25% | *p=0.0316* |
| | Polyploid, *ncd*$^{mut}$ | 47.36 ± 23.16 min | *p=0,4092 (ns)* | 1.97 ± 0.77 | *p=0,6650(ns)* | 0.0146 ± 0.0012 | *p<0.0001* | 88% | *p=0.0092* |

p = p-value : statistical significance related to the corresponding ploidy control.
ns = not significant
ND = not determined

### Supplementary Figure 2

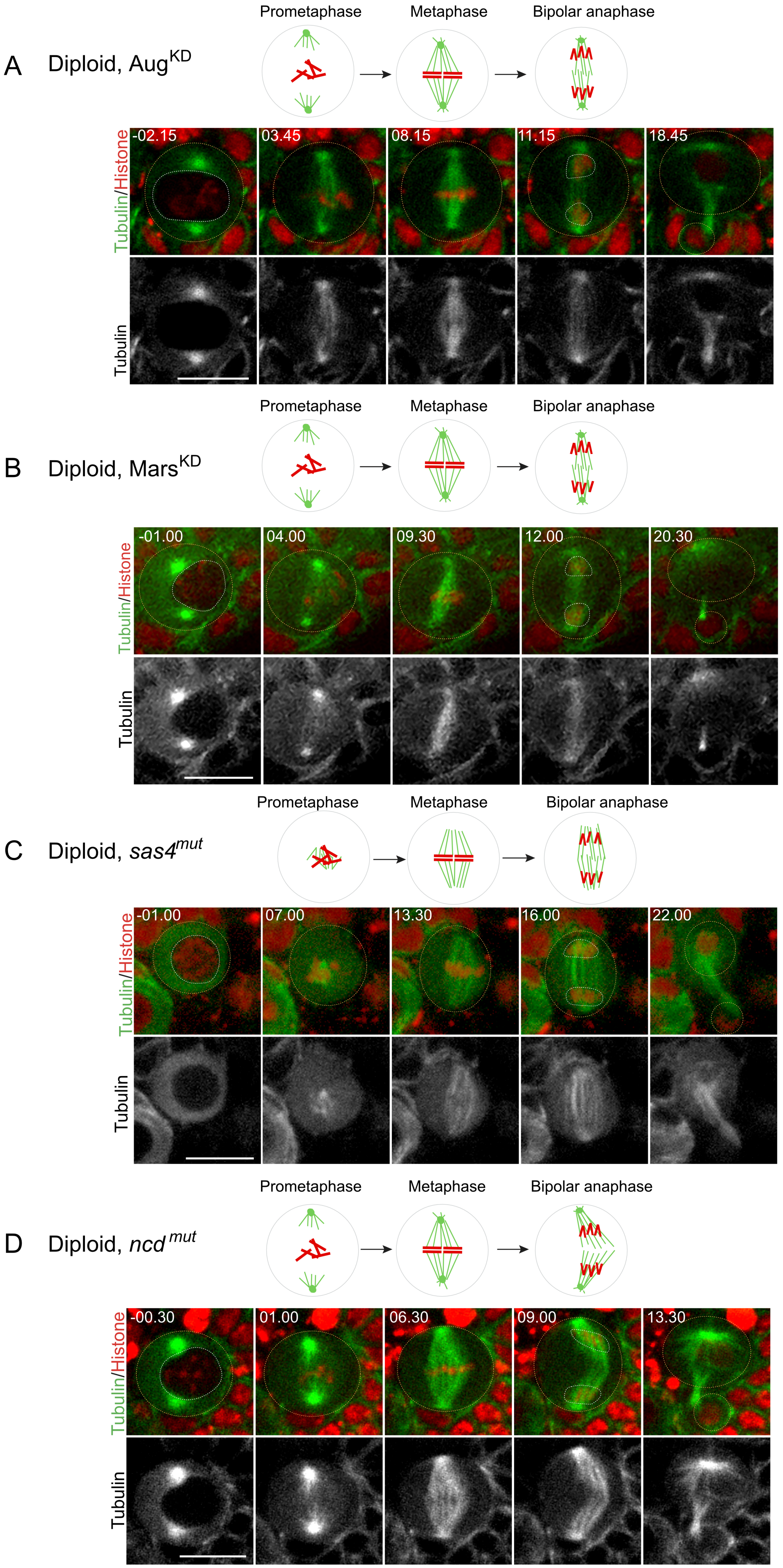

### Supplementary Figure 3

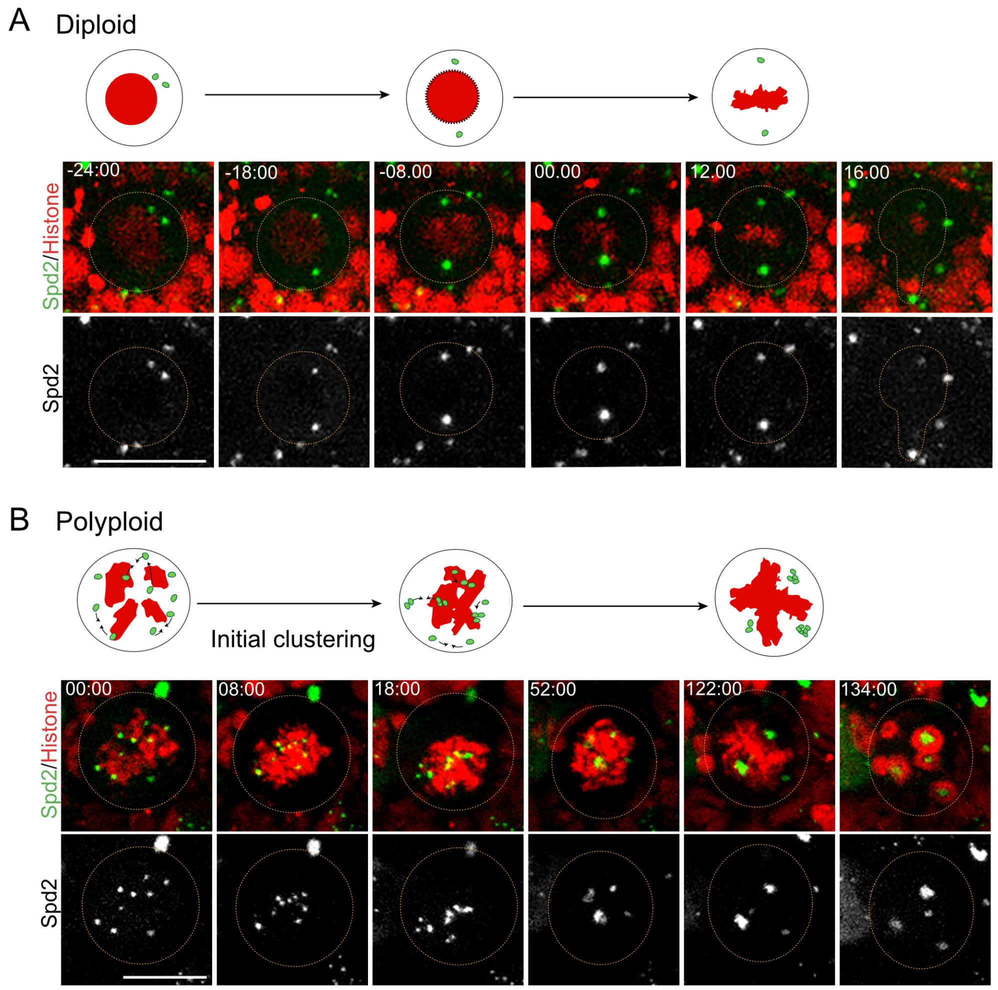

#### Supplementary Video legends

**Video 1:**

Ctrl mitotic NB expressing Tubulin-GFP (in green) and Histone 2B-RFP (in red). Time of mitosis is indicated in minutes(min).seconds(sec) and time 00.00 corresponds to nuclear envelope break down (NEBD). Still images from this video are shown in Figure 1A.

**Video 2:**

*Sqh*$^{mut}$ mitotic NB expressing Tubulin-GFP (in green) and Histone 2B-RFP (in red). Time of mitosis is indicated in hours(hr).min.sec and time 00.00.00 corresponds to the beginning of time-lapse acquisition. Still images from this video shown in Figure 1B.

**Video 3:**

Pav$^{KD}$ mitotic NB expressing Tubulin-GFP (in green) and Histone 2B-RFP (in red). Time of mitosis is indicated in hr.min.sec and time 00.00.00 corresponds to NEBD. Still images from this video shown in Figure 1C.

**Video 4:**

Small Pav$^{KD}$ mitotic NB expressing Tubulin-GFP (in green) and Histone 2B-RFP (in red). Time of mitosis is indicated in hr.min.sec and time 00.00.00 corresponds to NEBD. Still images from this video are shown in Supplementary Figure 1C.

**Video 5:**

Diploid, Aug$^{KD}$ mitotic NB expressing Tubulin-GFP (in green) and Histone 2B-RFP (in red). Time of mitosis is indicated in min.sec and time 00.00 corresponds to NEBD. Still images from this video are shown in Supplementary Figure 2A.

**Video 6:**

Polyploid, Aug$^{KD}$ mitotic NB expressing Tubulin-GFP (in green) and Histone 2B-RFP (in red). Time of mitosis is indicated in hr.min.sec and time 00.00.00 corresponds to NEBD. Still images from this video are shown in Figure 2B.

**Video 7:**

Polyploid, Mars$^{KD}$ mitotic NB expressing Tubulin-GFP (in green) and Histone 2B-RFP (in red). Time of mitosis is indicated in hr.min.sec and time 00.00.00 corresponds to NEBD. Still images from this video are shown in Figure 2F.

**Video 8:**

Diploid, Mars$^{KD}$ mitotic NB expressing Tubulin-GFP (in green) and Histone 2B-RFP (in red). Time of mitosis is indicated in min.sec and time 00.00 corresponds to NEBD. Still images from this video are shown in Supplementary Figure 2B.

**Video 9:**

Small polyploid, *sas4*$^{mut}$ mitotic NB expressing Tubulin-GFP (in green) and Histone 2B-RFP (in red). Time of mitosis is indicated in hr.min.sec and time 00.00.00 corresponds to NEBD. Still images from this video are shown in Figure 3A.

**Video 10:**

Diploid, *sas4*$^{mut}$ mitotic NB expressing Tubulin-GFP (in green) and Histone 2B-RFP (in red). Time of mitosis is indicated in min.sec and time 00.00 corresponds to NEBD. Still images from this video are shown in Supplementary Figure 2C.

**Video 11:**

Large polyploid, *sas4*$^{mut}$ mitotic NB expressing Tubulin-GFP (in green) and Histone 2B-RFP (in red). Time of mitosis is indicated in hr.min.sec and time 00.00.00 corresponds to NEBD. Still images from this video are shown in Figure 3B.

**Video 12:**

Polyploid, *ncd*$^{mut}$ mitotic NB expressing Tubulin-GFP (in green) and Histone 2B-RFP (in red). Time of mitosis is indicated in hr.min.sec and time 00.00.00 corresponds to NEBD. Still images from this video are shown in Figure 3F.

**Video 13:**

Diploid, *ncd*$^{mut}$ mitotic NB expressing Tubulin-GFP (in green) and Histone 2B-RFP (in red). Time of mitosis is indicated in min.sec and time 00.00 corresponds to NEBD. Still images from this video are shown in Supplementary Figure 2D.

**Video 14:**

Ctrl mitotic NB expressing Spd2-GFP (in green) and Histone 2B-RFP (in red). Time of mitosis is indicated in hr.min.sec and time 00.00.00 corresponds to NEBD Still images from this video are shown are shown in Supplementary Figure 3A.

**Video 15:**

$Pav^{KD}$ mitotic NB expressing Spd2-GFP (in green) and Histone 2B-RFP (in red). Time of mitosis is indicated in hr.min.sec and time 00.00.00 corresponds to NEBD. Still images from this video are shown in Supplementary Figure 3B.

**Video 16:**

Computational simulation of centrosome clustering (in green) with “metaphase-like plate” as DNA shape 1 (in red). Still images from this video are shown in Figure 4B.

**Video 17:**

Computational simulation of centrosome clustering (in green) with “3-pointed star” as DNA shape 2 (in red). Still images from this video are shown in Figure 4B.

**Video 18:**

Computational simulation of centrosome clustering (in green) with “4-pointed star” as DNA shape 3 (in red). Still images from this video are shown in Figure 4B.

**Video 19:**

Computational simulation of centrosome clustering (in green) with “5-pointed star” as DNA shape 4 (in red). Still images from this video are shown in Figure 4B.

**Video 20:**

Diploid, SakOE, $mad2^{mut}$ mitotic NB expressing Tubulin-GFP (in green) and Histone 2B-RFP (in red). Time of mitosis is indicated in min.sec and time 00.00 corresponds to NEBD. Still images from this video are shown in Figure 4D.

**Video 21:**

Diploid, SakOE, *mad2*$^{mut}$ mitotic NB expressing Tubulin-GFP (in green) and Histone 2B-RFP (in red). Time of mitosis is indicated in min.sec and time 00.00 corresponds to NEBD. Still images from this video are shown in Figure 4E.

#### Supplementary Materials

#### Supplementary Table 2:

| Parameter | | Description/reference |
|---|---|---|
| *Simulation* | | |
| Total time | 8 min | Mitosis duration was around 20 min for polyploid cells and clustering mostly happened in the first third of the time. |
| Viscosity | 1 pN. S/µm² | Cytoplasmic viscosity, as in [1] |
| *Microtubules* | | |
| Rigidity | 30 pN/µm² | Bending rigidity [2] |
| Polymerization speed | 0.26 µm/s | Growth speed during mitosis [3] |
| Depolymerization speed | -0.46 µm/s | Chosen to have a mean MT length of 3.5 µm [4] in a plausible range of values |
| Rescue rate | 0.22 /s | Chosen to have a mean MT length of 3.5 µm in a plausible range of values |
| Catastrophe rate | 0.2 /s | Chosen to have a mean MT length of 3.5 µm in a plausible range of values |
| Stall force | 5 pN | [5] |
| *Ncd* | | |
| Binding rate | 10 /s | Rate at which each head of Ncd can bind to a microtubule nearby. Chosen fast so that binding is not limiting |
| Binding range | 200 nm | Distance from which Ncd heads can bind to microtubules |
| Unbinding rate | 0.3 /s | Detachment of one head of Ncd dfrom the microtubule [6] (for conventional kinesin) |
| Unbinding force | 2.5 pN | Detachment force [7] (for kinesins) |
| Speed | -0.2 µm/s | Speed of motion of a bound motor domain ([8],[9]). |
| Stall force | 6 pN | Stall force of the motor domain of Ncd [7] |
| Spring stiffness | 100 pN/µm | Stiffness of the link between the two Ncd heads |
| *Centrosomes* | | |
| Viscosity | 200 pN s/µm² | Constrained the mobility of the centrosomes [1] |
| Radius | 0.5 µm | Radius of the centrosome objects |
| Number of MTs | 50 | Number of microtubules nucleated by each centrosome |
| *DNA surface* | | |
| Binding range | 0.2 µm | Distance from which microtubules can be caught by DNA entities placed on its surface. Bound to plus end of free microtubules (not caught yet) only. |

| | | |
|---|---|---|
| Binding rate | 5 /s | Fast binding |
| Unbinding rate | 0.01 /s | DNA surface entities stay fixed for a time on the microtubules plus end, preventing them from gliding on the surface |

#### Flies genetic crosses

Figure 1

Live imaging

Control: Tub-GFP, Hist-RFP / Cyo-GFP; ActGal4, Gal80$^{ts}$ / TM6,Tb

*Sqh*$^{mut}$ : *Sqh*$^{mut}$/FM7, Kruppel-GAL4, UAS-GFP ; Tub-GFP, Hist-RFP / Cyo-GFP

Fly crosses were maintained at 25°C

Pav$^{KD}$ : Pav$^{RNAi}$ / Cyo-GFP x Tub-GFP, Hist-RFP / Cyo-GFP; Actin-GAL4-GAL80$^{ts}$ / TM6,Tb

Crosses were maintained at 18°C after egg laying until reaching 1$^{st}$/2$^{nd}$ instar stages and then switched to 29°C for 24-48h.

Figure 2

Live imaging

Control : Aug$^{KD}$ : Tub-GFP,Hist-RFP / Cyo-GFP ; dgt2$^{RNAi}$ / TM6,Tb

x ActGal4-Gal80$^{ts}$ / TM6, Tb

Polyploid: Aug$^{KD}$ : Pav$^{RNAi}$ / Cyo-GFP; dgt2$^{RNAi}$ / TM6,Tb

x Tub-GFP,Hist-RFP / Cyo-GFP; ActGal4,Gal80ts/TM6,Tb

Control : Mars$^{KD}$ : Tub-GFP,Hist-RFP / Cyo-GFP ; mars$^{RNAi}$ / TM6,Tb

x ActGal4-Gal80$^{ts}$ / TM6, Tb

Polyploid, Mars$^{KD}$ : Pav$^{RNAi}$ / Cyo-GFP; mars$^{RNAi}$ / TM6,Tb

x Tub-GFP,Hist-RFP/Cyo-GFP; ActGal4-Gal80$^{ts}$ / TM6,Tb

Crosses were maintained at 18°C after egg laying until reaching 1$^{st}$/2$^{nd}$ instar stages and then switched to 29°C for 24-48h.

Figure 3

Live imaging

Control: *sas4*$^{mut}$ : Tub-GFP,Hist-RFP / Cyo-GFP ; *DSas4*$^{mut(S2214)}$ / TM6,Tb

Polyploid, *sas4*$^{mut}$ : Pav$^{RNAi}$,Hist-RFP / Cyo-GFP ; *DSas4*$^{mut(S2214)}$ / TM6,Tb x AsGal4,Tub-GFP / Cyo-GFP ; *DSas4*$^{mut(S2214)}$ / TM6,Tb

Control: *ncd*$^{mut}$ : Tub-GFP,Hist-RFP / Cyo-GFP ; *ncd*$^{1}$ / TM6,Tb

Polyploid, *ncd*$^{mut}$ : Pav$^{RNAi}$,Hist-RFP / Cyo-GFP ; *ncd*$^{1}$ / TM6,Tb

x AsGAL4,Tub-GFP / Cyo-GFP ; *ncd*$^{1}$ / TM6,Tb

Crosses were maintained at 25°C

Figure 4

Fixed analysis

Control : wt[118]

Polyploid : Pav$^{KD}$ : Pav$^{RNAi}$ / Cyo-GFP x Actin-GAL4-GAL80$^{ts}$ / Cyo-GFP

Crosses were maintained at 18°C after egg laying until reaching 1$^{st}$/2$^{nd}$ instar stages and then switched to 29°C for 24-48h.

Live imaging

SakOE, *mad2*$^{mut}$: pUbqSak-#68, *mad2*$^{mut}$/ TM6,Tb X Tub-GFP,Hist-RFP; *mad2*$^{mut}$

Supplementary Figure 1

Fixed analysis

Control: WT isogenic: WT flies were isogenized for the genetic background of *sqh* mutant, derived from WT female crossed with males FM7, Kruppel-GAL4, UAS-GFP)

*Sqh*$^{mut}$ : sqh$^{1}$/ FM7, Kruppel-GAL4, UAS-GFP

Fly lines were maintained at 25°C.

Pav$^{KD}$ : Pav$^{RNAi}$ / Cyo-GFP x Actin-GAL4-GAL80$^{ts}$ / Cyo-GFP

Crosses were maintained at 18°C after egg laying until reaching 1$^{st}$/2$^{nd}$ instar stages and then switched to 29°C for 24-48h.

Live imaging

Pav$^{KD}$ : Pav$^{RNAi}$ / Cyo-GFP x Tub-GFP, Hist-RFP / Cyo-GFP; Actin-GAL4-GAL80$^{ts}$ / TM6,Tb

Crosses were maintained at 18°C after egg laying and until reaching 1$^{st}$/2$^{nd}$ instar stages and then switched to 29°C for 24-48h.

Supplementary Figure 2

Live imaging

Diploid, $Aug^{KD}$ : Tub-GFP,Hist-RFP / Cyo-GFP ; $dgt2^{RNAi}$ / TM6,Tb x ActGal4-$Gal80^{ts}$ / TM6, Tb

Diploid, $Mars^{KD}$ : Tub-GFP,Hist-RFP / Cyo-GFP ; $mars^{RNAi}$ / TM6,Tb x ActGal4-$Gal80^{ts}$ / TM6, Tb

Crosses were maintained at 18°C after egg laying and until reaching $1^{st}/2^{nd}$ instar stages and then switched to 29°C for 24-48h.

Diploid, $sas4^{mut}$ : Tub-GFP,Hist-RFP / Cyo-GFP ; $sas4^{mut}$ / TM6,Tb

Diploid, $ncd^{mut}$ : Tub-GFP,Hist-RFP / Cyo-GFP ; $ncd^{mut}$ / TM6,Tb

Crosses were maintained at 25°C.

Supplementary Figure 3

Live imaging

Diploid : Spd2-GFP,Hist-GFP / Cyo-GFP

Polyploid : $Pav^{RNAi}$ / Cyo-GFP x Spd2-GFP,Hist-RFP / Cyo-GFP ; ActGal44,$Gal80^{ts}$ / TM6,Tb

Crosses were maintained at 18°C after egg laying until reaching $1^{st}/2^{nd}$ instar stages and then switched to 29°C for 24-48h.

#### Supplementary References


1. Letort, G., Nedelec, F., Blanchoin, L., and Thery, M. (2016). Centrosome centering and decentering by microtubule network rearrangement. Molecular biology of the cell *27*, 2833-2843.
2. Gittes, F., Mickey, B., Nettleton, J., and Howard, J. (1993). Flexural rigidity of microtubules and actin filaments measured from thermal fluctuations in shape. The Journal of cell biology *120*, 923-934.
3. Gallaud, E., Caous, R., Pascal, A., Bazile, F., Gagne, J.P., Huet, S., Poirier, G.G., Chretien, D., Richard-Parpaillon, L., and Giet, R. (2014). Ensconsin/Map7 promotes microtubule growth and centrosome separation in Drosophila neural stem cells. The Journal of cell biology *204*, 1111-1121.
4. Rhys, A.D., Monteiro, P., Smith, C., Vaghela, M., Arnandis, T., Kato, T., Leitinger, B., Sahai, E., McAinsh, A., Charras, G., et al. (2018). Loss of E-cadherin provides tolerance to centrosome amplification in epithelial cancer cells. The Journal of cell biology *217*, 195-209.
5. Dogterom, M., and Yurke, B. (1997). Measurement of the force-velocity relation for growing microtubules. Science (New York, N.Y.) *278*, 856-860.
6. Howard, J., Hudspeth, A.J., and Vale, R.D. (1989). Movement of microtubules by single kinesin molecules. Nature *342*, 154-158.
7. Rupp, B., and Nedelec, F. (2012). Patterns of molecular motors that guide and sort filaments. Lab on a chip *12*, 4903-4910.
8. Oladipo, A., Cowan, A., and Rodionov, V. (2007). Microtubule motor Ncd induces sliding of microtubules in vivo. Molecular biology of the cell *18*, 3601-3606.
9. Fink, G., Hajdo, L., Skowronek, K.J., Reuther, C., Kasprzak, A.A., and Diez, S. (2009). The mitotic kinesin-14 Ncd drives directional microtubule-microtubule sliding. Nature cell biology *11*, 717-723.
